# High-throughput single-cell functional elucidation of neurodevelopmental disease-associated genes reveals convergent mechanisms altering neuronal differentiation

**DOI:** 10.1101/862680

**Authors:** Matthew A. Lalli, Denis Avey, Joseph D. Dougherty, Jeffrey Milbrandt, Robi D. Mitra

## Abstract

The overwhelming success of exome- and genome-wide association studies in discovering thousands of disease-associated genes necessitates novel high-throughput functional genomics approaches to elucidate the mechanisms of these genes. Here, we have coupled multiplexed repression of neurodevelopmental disease-associated genes to single-cell transcriptional profiling in differentiating human neurons to rapidly assay the functions of multiple genes in a disease-relevant context, assess potentially convergent mechanisms, and prioritize genes for specific functional assays. For a set of 13 autism spectrum disorder (ASD) associated genes, we demonstrate that this approach generated important mechanistic insights, revealing two functionally convergent modules of ASD genes: one that delays neuron differentiation and one that accelerates it. Five genes that delay neuron differentiation (*ADNP, ARID1B, ASH1L, CHD2*, and *DYRK1A*) mechanistically converge, as they all dysregulate genes involved in cell-cycle control and progenitor cell proliferation. Live-cell imaging after individual ASD gene repression validated this functional module, confirming that these genes reduce neural progenitor cell proliferation and neurite growth. Finally, these functionally convergent ASD gene modules predicted shared clinical phenotypes among individuals with mutations in these genes. Altogether these results demonstrate the utility of a novel and simple approach for the rapid functional elucidation of neurodevelopmental disease-associated genes.

## Introduction

The tremendous progress in identifying disease-associated genes and variants has far outpaced the discovery of the functions and pathological mechanisms of these genes. Exome- and genome-wide sequencing studies have identified ∼5500 single gene disorders and traits caused by mutations in over 3800 genes^1^. Over 1100 of these genes have been causally linked to neurodevelopmental disorders^2^. In autism spectrum disorder (ASD) alone, recent exome sequencing studies have identified over 100 genes that cause ASD when a single copy is mutated to a loss-of-function allele^3–6^. This genetic heterogeneity provides a substantial challenge to the development of broadly useful therapeutics. If, at an extreme, each disease-associated gene follows a separate mechanistic route, then each will require the development of an independent therapeutic. On the other hand, if subsets of these genes converge in their mechanisms, then these points of convergence would be logical targets for more broadly applicable therapeutics that apply to the entire subset. Identifying convergent mechanisms across diverse disease-associated genes first requires establishing a high-throughput and disease-relevant model system to both perturb numerous genes and systematically assess the functional consequences.

While animal models and patient-derived induced pluripotent stem cell (iPSC) models are powerful tools for the study of disease mechanisms, these systems are generally low-throughput, require long generation times, and results can vary across laboratories, strains, or individuals^7, 8^. A rapid, reproducible, and disease-relevant system in which multiple genes could be studied in parallel would fill an important gap in the functional genomics toolbox and enable direct comparison across genes to assess their mechanistic convergence. Fortunately, recent technological advancements coupling CRISPR/Cas9 transcriptional repression to single-cell RNA sequencing (scRNA-seq) enable high-throughput perturbation of multiple genes in a single batch with a parallel functional readout of the transcriptional consequences^9–12^. Such an approach holds great promise for efficiently defining the functional consequences of dominant loss-of-function mutations, as transcriptional repression can phenocopy haploinsufficiency. As the pathology of many neurodevelopmental diseases likely arises during neural development, especially when proliferating progenitors are differentiating into post-mitotic neurons^13^, we set out to establish a scalable functional genomics approach in a simple human cellular model of neuron differentiation. Further, as a large fraction of causative genes in ASD are haplo-insufficient transcriptional regulators^4, 6, 14^, we tested this approach on a select subset of such genes to determine what insights into pathological mechanisms can be gleaned by measuring the transcriptional consequences of their perturbation.

Here, we used catalytically inactive Cas9-based transcriptional repression (dCas9-KRAB) to knock down the expression of 13 ASD-related genes in a human cellular model of neuronal differentiation and captured the resulting transcriptional consequences using scRNA-seq. For all candidate genes, we identified individual transcriptional signatures after repression. We found that many ASD genes altered the trajectory of neuronal differentiation when repressed. Furthermore, by clustering the disease genes by their shared transcriptional changes after perturbation, we identified sets of diverse autism genes that acted either by delaying or accelerating neuronal differentiation. Transcriptional convergence of these genes at the pathway level generated specific mechanistic hypotheses which we then tested and confirmed by combining individual knock-down experiments with live-cell imaging. Supporting the validity of our model of neuronal differentiation, we replicated our key results in an orthogonal system using iPSC-derived neural progenitor cells. Finally, we show that clustering of ASD genes by experimental data predicted shared clinical phenotypes of individuals with mutations in these genes. These results demonstrate the promise of a high-throughput functional genomics screening platform to identify convergently disrupted cellular pathways across diverse causative genes in a simple cellular model of neuronal differentiation.

## Results

### Establishing a human model of neuronal differentiation for high-throughput disease gene perturbation

Human neuronal models are needed for studying neurodevelopmental disorders such as ASD^8^. While human iPSC-derived neurons are a powerful cellular model system, the genetic heterogeneity, variability of neuronal differentiation, and technical difficulties achieving efficient transcriptional modulation in these cells complicate multiplexed transcriptional and phenotypic analyses^15, 16^. Therefore, we aimed to establish a tractable, diploid human neuronal model amenable to differentiation and transcriptional perturbation to enable high-throughput evaluation of the consequences of disease-associated gene repression. We selected the LUHMES neural progenitor cell line as such a model for their ease of use, capacity for rapid differentiation into post-mitotic neurons, and suitability for high-content imaging^17–20^. Recent studies have used LUHMES to model neurodevelopmental disorders and their underlying pathways^21, 22^. To further validate the relevance of these cells, we performed RNA sequencing (RNA-seq) analysis of LUHMES cells at multiple time points after inducing differentiation. Hierarchical clustering analysis of differentially expressed genes across the differentiation time-course confirmed that differentiation of LUHMES was rapid and reproducible (Pearson’s r^2^ between replicates > 0.99), with biological replicates clustering together and samples arranged temporally by their day of differentiation (Figure 1a, Supplementary Figure 1A). Genes that were down-regulated during differentiation were enriched for cell-cycle markers such as *CCND2*, genes involved in proliferation (*MKI67* and *TP53*), as well as the canonical neural stem cell marker gene *SOX2*. Genes that increased expression during differentiation included known neuronal markers *MAP2* and *DCX*, and were heavily enriched for critical neurodevelopmental pathways including axon growth, synaptic development, and neuron migration (Fig 1b Left). Importantly, genes expressed during differentiation were strongly enriched for genes implicated in a variety of neurological disorders, including schizophrenia, bipolar disorder, and ASD (Fig 1b Right).

**Figure 1:**
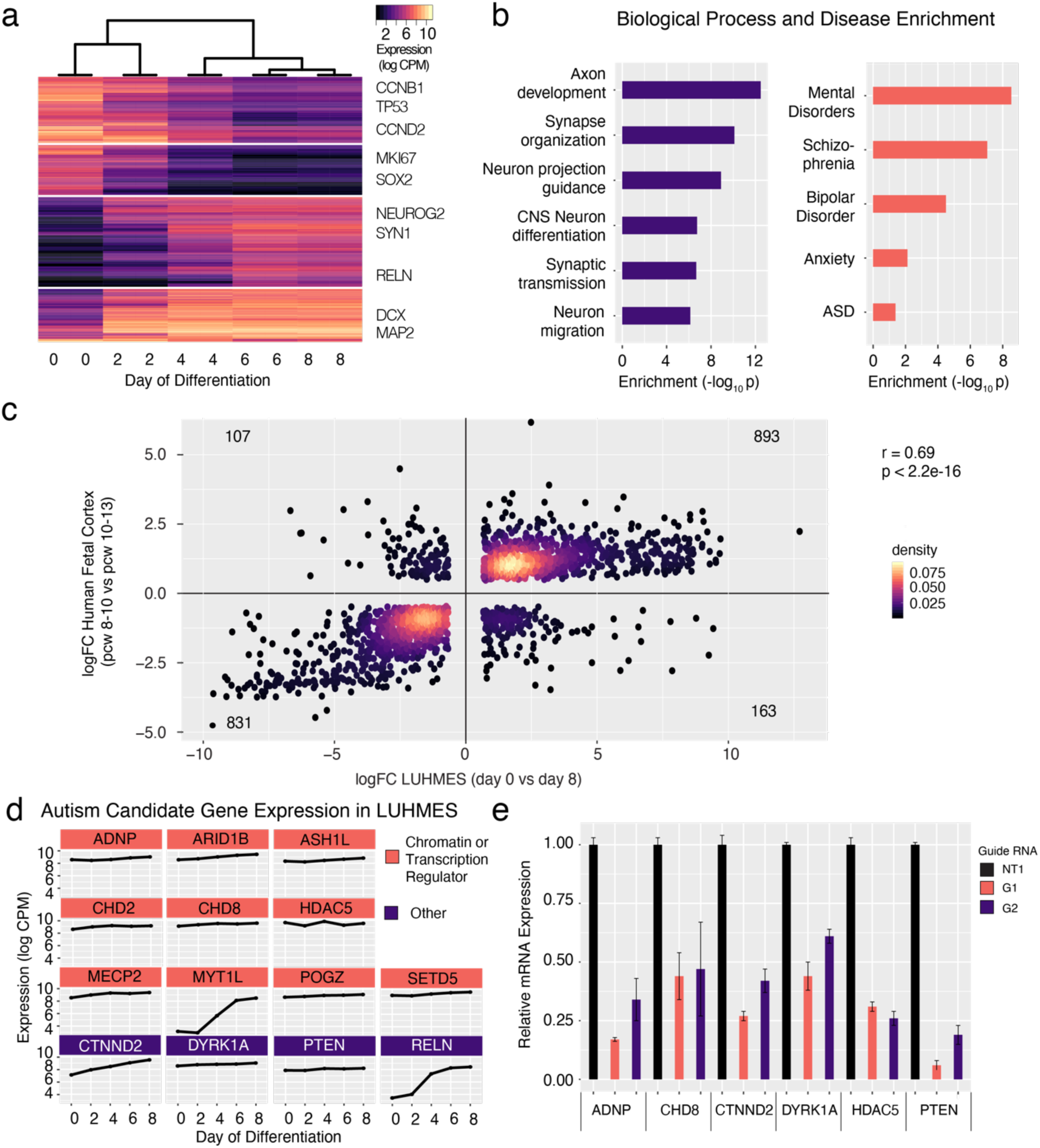
LUHMES are a tractable, disease-relevant model of human neuronal differentiation amenable to perturbation. **a)** Hierarchical clustering of bulk RNA-seq time-course expression data indicates rapid and reproducible neuronal differentiation. Two replicates for each timepoint were performed. **b**) Genes induced during LUHMES differentiation are enriched for relevant biological processes (Left) and neurological disorders (Right). **c)** Differentially expressed genes during LUHMES differentiation are highly correlated with transcriptional changes during early human fetal corticogenesis. **d**) High-confidence autism-causing genes, selected for perturbation experiments, are highly expressed at baseline or are increasingly expressed in LUHMES during differentiation and were selected for roles in transcriptional regulation. **e**) Efficient dCas9-KRAB repression of individual target genes using the designated guide RNAs. n = 3 biological replicates for all qPCR experiments. Values represent mean ± SEM. NT1: Nontargeting control guide RNA. G1: gRNA 1, G2: gRNA 2. pcw: post-conception week.

Despite being a mesencephalic-derived neuronal progenitor line best characterized for its ability to differentiate into dopaminergic neurons, cell type-specific expression analysis (CSEA) of differentiated LUHMES revealed that these neurons have transcriptional profiles that are highly similar to a range of neuronal subtypes relevant to neurological disorders (Supp Figure 1B)^23^. Specifically, transcriptomes of differentiated cells resembled striatal dopaminergic neurons as expected but also matched some cortical, forebrain, and spinal cord neuron types. Next, to assess the extent to which *in vitro* differentiation of LUHMES cells captures aspects of human brain development, we performed a transition-mapping approach comparing differentially expressed genes during LUHMES differentiation to the BrainSpan Atlas of Developing Human Brain (Methods)^24, 25^. We found that changes in gene expression during *in vitro* differentiation closely mirror transcriptional differences that occur in the early developing human fetal neocortex (Pearson’s r = 0.69, Fig 1c). This strong overlap suggests that LUHMES differentiation faithfully recapitulates many of the transcriptional pathways that are utilized during this critical neurodevelopmental window. Since LUHMES *in vitro* differentiation produces only a single neuronal cell type, some important disease-associated phenomena such as shifts in neuronal cell fate decisions or aberrations in region-specific gene regulatory networks will not be captured by this system. However, as core transcriptional programs that control neuronal differentiation and maturation are largely conserved across neuronal subtypes^26^, we can model these critical disease-relevant processes using a simple *in vitro* system.

To establish that LUHMES is an appropriate model specifically for the study of ASD genes, we analyzed the 25 highest-confidence autism-causing genes in the SFARI database (category 1), a manually curated database of ASD-associated genes^27^. We found that 22/25 (88%) were highly expressed in these cells across differentiation time points. We selected 11 of these genes, plus 2 additional syndromic ASD genes (*CTNND2* and *MECP2*)^28, 29^ for perturbation experiments (Table 1, Fig 1d). *HDAC5* was included as a non-associated gene which is highly expressed in neuronal progenitors where it may regulate stem cell proliferation^30^. Genes were selected for their roles in transcriptional regulation (10/14), and because they are highly likely to act through haploinsufficiency^31, 32^ (Table 1). Although many of these genes are co-expressed during neurodevelopment, module assignment of these genes by integrative bioinformatics approaches has not enabled specific mechanistic predictions about the potential convergence of their molecular targets^26, 33^. We expect this set of genes to be broadly representative of transcriptional regulators implicated in neurodevelopmental disorders and well-suited to demonstrate the feasibility of our approach.

**Table 1:**
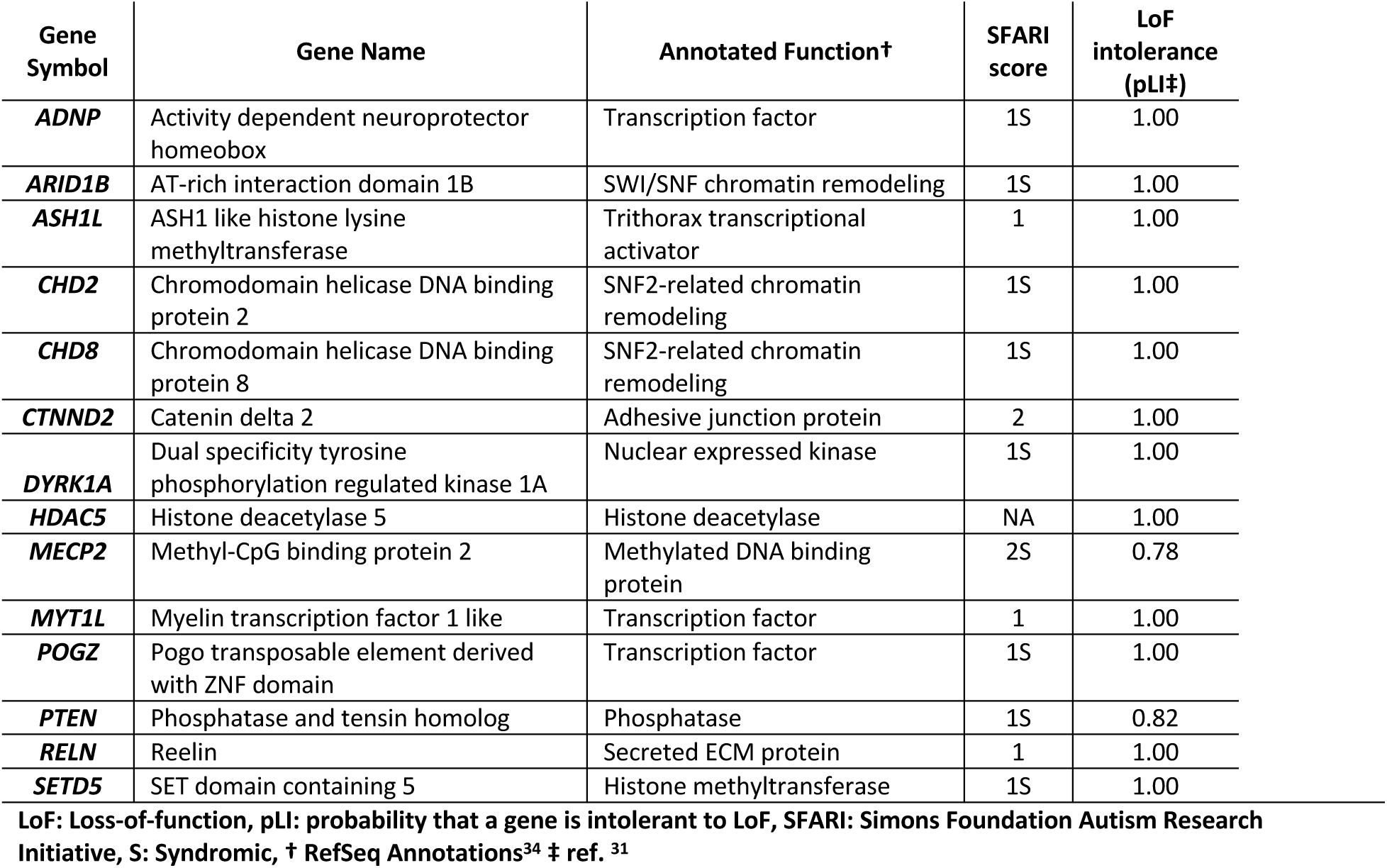
Description of candidate genes selected for perturbation experiments.

We next sought to determine whether the expression of candidate genes could be efficiently knocked down in LUHMES cells, a prerequisite for perturbation assays. Three guide RNAs (gRNAs) per candidate gene were cloned into a CRISPR-repression optimized vector that also allows recovery of the gRNA from scRNA-seq ^35–37^. We validated the efficacy of repression for 2 gRNAs targeting each of 6 candidate genes using quantitative real-time PCR (qPCR) in LUHMES neural progenitor cells constitutively expressing dCas9-KRAB. All tested gRNAs induced significant downregulation of their target gene, with 11/12 eliciting a knockdown greater than 50% (Fig 1e), a level that should phenocopy the autosomal-dominant loss-of-function modes of our candidate genes. Altogether, these data support LUHMES as a relevant and facile cellular model to evaluate the downstream consequences of transcriptional perturbation of neurodevelopmental genes.

### Pooled repression of ASD genes and scRNA-seq

We produced a lentivirus pool that contained vectors expressing gRNAs targeting all 14 candidate genes (3 gRNAs per gene), along with 5 non-targeting control gRNA sequences, for a total of 47 gRNAs. Given the high success rate of gene knockdown in dCas9-KRAB LUHMES by all tested gRNAs, and to enable high-scale perturbation screening experiments, we did not validate the repression efficiency of all gRNAs individually. We infected dCas9-KRAB expressing LUHMES neuronal progenitors at a low multiplicity of infection such that most cells received 0 or 1 gRNAs according to a Poisson distribution. Cells infected by a gRNA-expressing lentivirus were selected by growth in media containing puromycin for 4 days, then the cells were induced to differentiate according to published protocols ^17^(Methods). To allow sufficient time for CRISPR repression, we differentiated LUHMES for 7 days, a timepoint when differentiation appeared largely complete by RNA-seq. We then profiled the transcriptomes of more than 14,000 cells at this timepoint using droplet-based scRNA-seq across two replicate experiments^38^ (Figure 2a,b).

**Figure 2:**
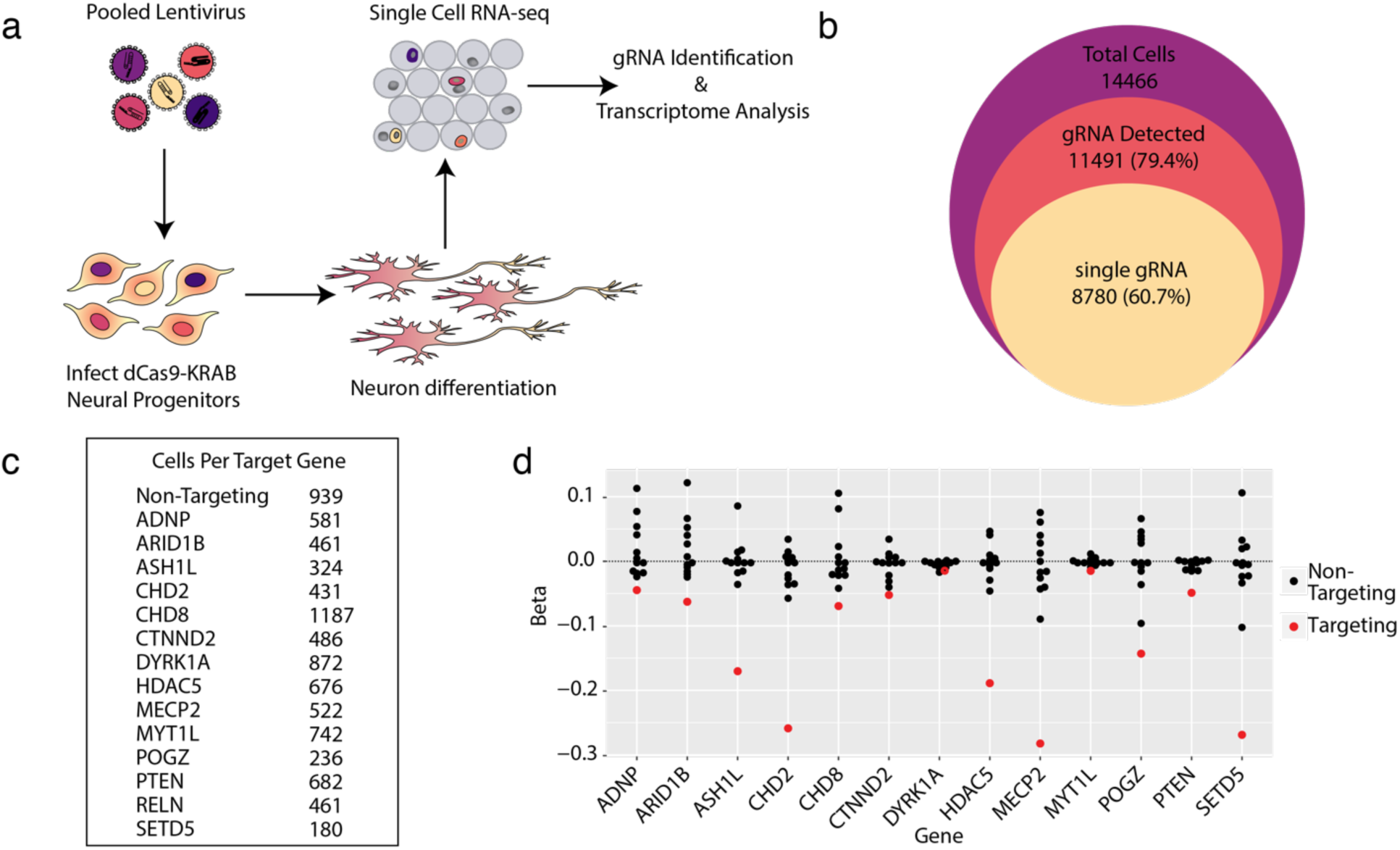
Single-cell RNA-sequencing is an efficient readout of multiplexed gene repression in a human model of neuronal differentiation. **a)** Schematic of pooled repression of ASD genes in LUHMES. **b**) The number of total single cells that were collected (purple area), that express at least one gRNA (red), and express a single unique gRNA (yellow) demonstrates the efficient recovery of gRNAs from single-cell RNA-seq data. **c)** Hundreds of cells targeting all 14 genes were recovered, as well as 939 cells with non-targeting gRNAs. **d)** MIMOSCA beta coefficients (arbitrary units) for targeting and non-targeting guides are shown for each targeted gene. Three targeting gRNAs for each gene are merged. Beta < 0 represents repression. In all cases, targeting gRNAs have negative beta coefficients on target gene expression.

Using a specific gRNA enrichment PCR, we were able to detect gRNA expression for the vast majority (∼80%) of cells and restricted our analysis to the 8780 high-quality cells with only a single gRNA to ensure only one perturbation per cell (Fig 2b). Cells with gRNAs targeting all candidate genes, and non-targeting controls, were represented in this dataset (Fig 2c). To evaluate the efficacy of ASD-gene repression in the pooled experiment, we grouped single cells by their detected gRNAs and visualized expression of all targeted genes across these groups (Supplementary Figure 2a)^39, 40^. This analysis revealed efficient on-target repression for 13/14 genes in our library. The number of reads for *RELN* was too low in single-cell data to evaluate efficiency of repression due to its low expression level. We used the MIMOSCA pipeline to further evaluate the knock-down efficiency of gRNAs on their target genes, which confirmed strong on-target repression^11^ (Fig 2d). Almost all individual gRNAs elicited significant repression of their target genes (Sup Fig 2B-D), and estimated knock-down efficiencies were highly reproducible between replicate experiments (Sup Fig 2E).

### ASD-gene repression alters trajectory of neuronal differentiation

The efficient repression of targeted genes in the pooled experiment demonstrated above led us to assess the unique and shared downstream consequences of ASD-gene repression in human neurons. Specifically, we wanted to directly test the hypothesis that some of the ASD genes might alter the dynamics of neuronal differentiation. To this end, we reconstructed a pseudo-temporal trajectory reflecting gene expression changes in our dataset and projected cells onto this pseudotime path (Figure 3a)^41^. Recent single-cell CRISPR experiments have shown the advantages of trajectory analysis over global clustering based approaches, which can be insensitive to detecting more subtle phenotypes in pooled experiments (Supplementary Figure 3, Methods)^42–44^. Importantly, as all cells were differentiated for 7 days, more than 99% of single cells were post-mitotic as assessed by the absence of proliferation markers *MKI67* and *TOP2A*. However, pseudotime analysis indicated heterogeneity in the progression of differentiation at the single-cell level. Two neuronal marker genes (*MAP2* and *DCX*) showed a gradual increase in expression across pseudotime. In contrast, two genes known to be important for neural progenitor cell proliferation (*TP53* and *CDK4*) showed a rapid drop in expression across pseudotime. These observations suggested the axis of pseudotime corresponds to the progression of neuronal differentiation (Fig 3b). Consistent with this notion, these four genes exhibit a similar pattern of expression over a time-course of LUHMES differentiation (Fig 3c). To further examine the relationship between pseudotime and neuronal differentiation, we identified the marker genes for each pseudotime state and plotted their expression across the differentiation time-course RNA-seq dataset. This analysis showed that marker genes of early pseudotime states (1-3) are highly expressed in early neuron differentiation (days 0-4) (Supplementary Figure 4A-C). Marker genes of later pseudotime states (4-6) are highly expressed during later neuron differentiation (days 4-8) (Sup Figure 4D-F). Altogether, these data support the interpretation of pseudotime as an axis corresponding to the progression of neuronal differentiation after cells have completed their final division.

**Figure 3:**
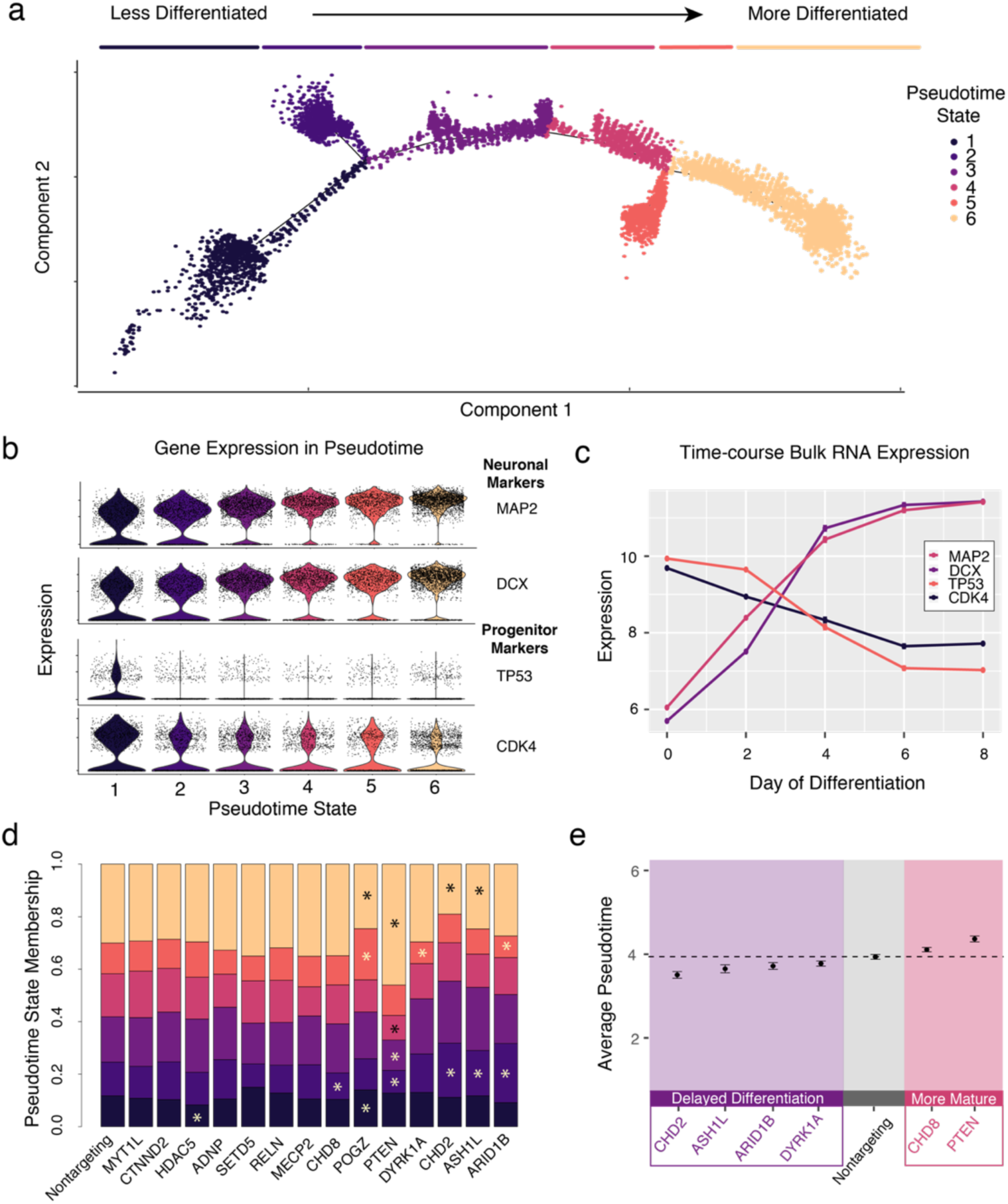
Pseudotime analysis reveals ASD-gene repression induced alterations in differentiation trajectory and modules of ASD-genes delaying or accelerating neuronal differentiation. **a**) Pseudotime ordering of all single cells reveals a continuous trajectory of cell states corresponding to neuronal differentiation. Line segments along the trajectory are called ‘Pseudotime States’ and cells are colored by these states. **b**) Neuronal markers (MAP2 and DCX) increase along the pseudotime trajectory, while progenitor markers (TP53 and CDK4) decrease. **c**) These marker genes exhibit correlated patterns in time-course bulk RNA expression. n = 2 replicates for each timepoint. Expression values = mean ± SEM. **d**) Repression of some genes alter pseudotime state membership proportions. Significant enrichments and depletions are marked with asterisks (chi-squared test, p < 0.01). **e**) Significantly decreased or increased average pseudotime scores relative to cells with non-targeting gRNAs (t-test, p < 0.01), indicate delayed or accelerated neuronal maturation. Boxed in purple is a set of ASD genes that delay neuronal differentiation by this metric. Boxed in red is a pair of genes, *CHD8* and *PTEN*, that promote neuronal maturation.

To test whether any candidate gene knockdown shifted the developmental trajectory of the differentiating neurons, we computed the proportions of cells in each pseudotime state for each knockdown condition. Indeed, we found that several perturbations significantly altered the proportions of cells in specific pseudotime states. Specific enrichment or depletion of pseudotime state clusters for each target gene are shown in Fig 3d (chi-squared tests, *p* < 0.01). As an indicator of differentiation status, we next computed the average value of pseudotime across all cells in each knockdown condition. By this metric, we found that four of the ASD genes (*CHD2*, *ASH1L*, *ARID1B*, and *DYRK1A*) delayed neuronal differentiation when repressed whereas two genes (*PTEN* and *CHD8*) accelerated neuronal differentiation (Fig 3e, p < 0.01, t-test). These results demonstrate the utility of pseudotime analysis to investigate whether subsets of disease-associated genes alter the progression of neuronal differentiation in a pooled perturbation experiment.

### Recurrently dysregulated genes highlight convergent mechanisms of ASD genes

We noticed that specific pseudotime state enrichment or depletion was not perfectly shared among the sets of genes that accelerated or delayed differentiation (Fig 3d). This raises the possibility that although groups of genes may act similarly to promote (or delay) neuron differentiation, they may do so through different molecular mechanisms. We therefore sought to further dissect the transcriptional networks affected by gene perturbation using differential gene expression analysis to learn whether these networks converge across sets of ASD genes. Due to the low number of cells available for analysis of *SETD5* and *POGZ* (Fig 2c), we excluded these cells from differential expression analyses. For each of the remaining genes, we found dozens to hundreds of differentially expressed genes for each knock-down (Supplementary Table 1).

To identify potential transcriptional convergence of diverse ASD-causing genes, we grouped cells by targeted gene then clustered their transcriptional profiles using only genes that were found to be differentially expressed across three or more ASD-gene knockdowns. The grouping of ASD genes via hierarchical clustering largely recapitulated the results of pseudotime analysis (Supplementary Figure 5A). Gene Ontology analysis of the set of dysregulated genes showed an enrichment of neuronal differentiation terms in the perturbed transcriptomes, supporting the interpretation that the misregulation of these ASD-associated genes alters neuronal differentiation (Sup Fig 5B). This analysis identified *ADNP* as another gene delaying neuronal differentiation and indicates that the repression of *ADNP, ASH1L, CHD2*, and *DYRK1A* delays neuronal differentiation through a shared transcriptional pathway. However, since our earlier pseudotime analysis revealed heterogeneity in the progression of neuronal differentiation amongst these genes, the shared transcriptional changes that we observed in our differential expression analysis may have been caused by distinct transcriptional changes that occurred early in neuronal differentiation. To address this possibility, we leveraged the power of single-cell data to explicitly account for differences in neuronal maturity by stratifying samples based on pseudotime (Sup Fig 5C, Methods). In order to retain enough cells per group required for differential gene expression analysis^45^, we dichotomized pseudotime status as either ‘early’ or ‘late’ and then recomputed differentially expressed genes for each ASD-gene knockdown within each group (Supplementary Tables 2 and 3).

We then clustered pseudotime-stratified samples based on recurrently dysregulated genes (*i.e.* genes that were differentially expressed in three or more ASD-gene knockdown samples) and found shared and distinct patterns of transcriptional dysregulation (Figure 4a). This analysis grouped samples first by early and late pseudotime status, and then into stage-specific subsets of ASD genes. Because genes that acted to delay differentiation clustered together at the early stage, we can infer that the delayed neuronal maturation detected in our pseudotime analysis is a consequence of early-stage transcriptional dysregulation. Furthermore, the sets of transcriptional targets of these genes shared significant overlap (all pair-wise hypergeometric p-values < 10^-22^, Fig 4b), implying that these genes act through a convergent regulatory pathway that functions early in neuronal maturation. This analysis also provided information about the regulatory hierarchy of these genes, with *CHD2* downregulated by *ADNP* or *ARID1B* repression, and *ASH1L* downregulated by *ARID1B* repression. Despite the cells being post-mitotic, Gene Ontology enrichment of the recurrently dysregulated genes in the early stage samples highlighted specifically disrupted processes, namely the G2/M transition of cell cycle and negative regulation of cell development (Fig 4c). The disrupted genes themselves are not core cell-cycle regulators so this signature may rather reflect cell-cycle disruptions that occurred earlier in the differentiation protocol (Supplementary Table 4). Together these results suggest that *CHD2, ASH1L, ARID1B, DYRK1A*, and *ADNP* comprise a convergent functional module of ASD genes acting on a shared gene regulatory pathway active in early neurodevelopment, and that their haploinsufficiency causes cell-cycle disruption and impedes neuronal differentiation.

**Figure 4:**
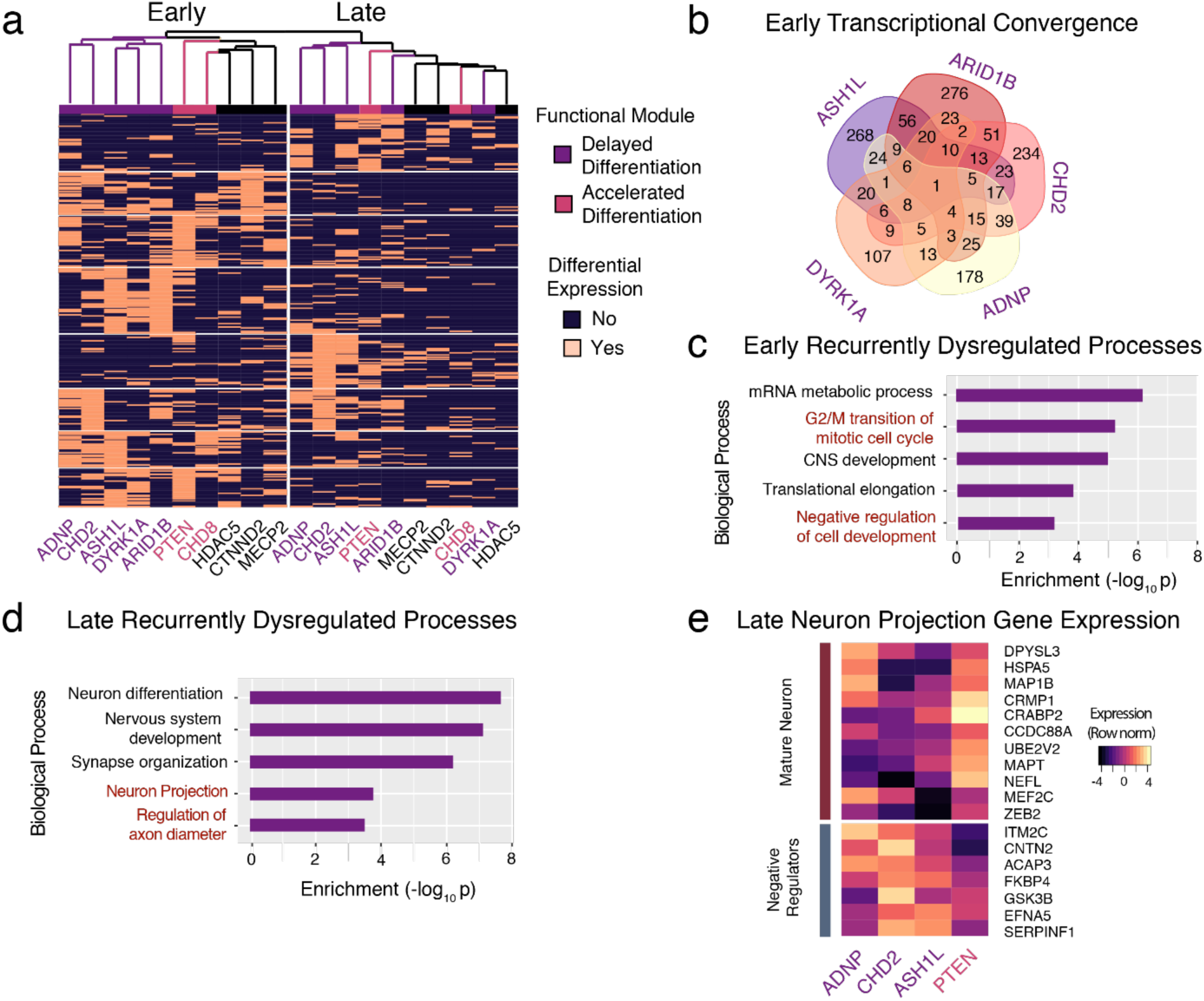
Single-cell differential gene expression analysis identifies early and late stage convergent modules of ASD genes. **a**) Hierarchical clustering of ASD knockdown profiles using genes differentially expressed in 3 or more ASD-candidate perturbations reveal a convergence of the transcriptional pathways dysregulated at early and late stages. At the early stage (left column), *ADNP, CHD2, ASH1L, DYRK1A*, and *ARID1B* form a transcriptionally convergent module of ‘Delayed Differentiation’ (purple). At the late stage (right column), *ADNP, CHD2*, and *ASH1L* continue to converge. **b**) Venn diagram shows significant overlap of differentially expressed genes across 5 ASD-genes at the early stage. All pairwise overlaps have p-values < 10^-22^ by hypergeometric testing. **c)** Gene Ontology enrichment analysis of early-stage recurrently dysregulated genes highlights relevant biological processes disrupted and predicts disrupted G2/M transition and cellular maturation for ‘Delayed Differentiation’ genes. **d**) Enrichment analysis of late-stage recurrently dysregulated genes highlights relevant biological processes disrupted. **e**) Expression of neuron projection genes (Gene Ontology 0010975) in the late stage samples predicts disrupted neurite extension for *PTEN* but an enhanced phenotype for *ADNP, CHD2*, and *ASH1L*.

Notably, *ADNP, CHD2*, and *ASH1L* were also clustered in late stage cells implying they continue to share downstream molecular targets in maturing neurons. In contrast, hierarchical clustering did not support a convergence of the genes that accelerate differentiation, *PTEN* and *CHD8*, at either the early or late stages. We found that the recurrently dysregulated genes among late stage samples were enriched for processes specific to neuron maturation such as synapse organization, neuron projection, and regulation of axon diameter (Fig 4d). The specificity of these terms highlights the added molecular resolution we achieved by accounting for differences in pseudotime in our analysis.

We next sought to generate specific predictions regarding the effects of ASD-gene repression on neuronal projections (axons and dendrites). Therefore, we clustered samples by the expression of the dysregulated neuron projection genes and found that the clustering of *ADNP, CHD2*, and *ASH1L* was driven by the decreased expression of neuron maturation markers such as *MAPT, NEFL*, and *MAP1B* with concomitant up-regulation of annotated negative regulators of neuron projection and differentiation (Fig 4e). These genes were driven in the opposite direction by *PTEN*. From these results, we predicted that knockdown of *ADNP, CHD2*, and *ASH1L* would decrease the outgrowth of neuronal projections, whereas knockdown of *PTEN* would enhance this process.

### Live-cell imaging reveals abnormalities in proliferation and neurite extension and confirms transcriptome-based predictions

Our single-cell transcriptional analyses allowed us to make several explicit predictions about the consequences of ASD-gene repression on cellular phenotypes. Specifically, we predicted that if *ADNP, ARID1B, ASH1L, CHD2*, or *DYRK1A* are repressed, then we would observe a reduction in neural progenitor cell proliferation. In contrast, we expected *PTEN* repression to promote proliferation. For neurite extension in differentiating neurons, we predicted that repression of *ADNP, CHD2*, and *ASH1L* would decrease outgrowth, whereas repression of *PTEN* would enhance extension. To test these predictions, we implemented live-cell imaging to measure cellular proliferation and neurite extension after individual knock-down of ASD genes (Figure 5a). We produced lentivirus expressing gRNAs that target candidate ASD genes and used them to infect dCas9-KRAB neuronal progenitor cells in an arrayed format. We imaged cells under both proliferative and differentiative conditions every 4 hours for 3 or 5 days, respectively, using the IncuCyte live-cell imaging system. Representative images for proliferation and neurite extension are shown (Fig 5b-c, Supplementary Figure 6).

**Figure 5:**
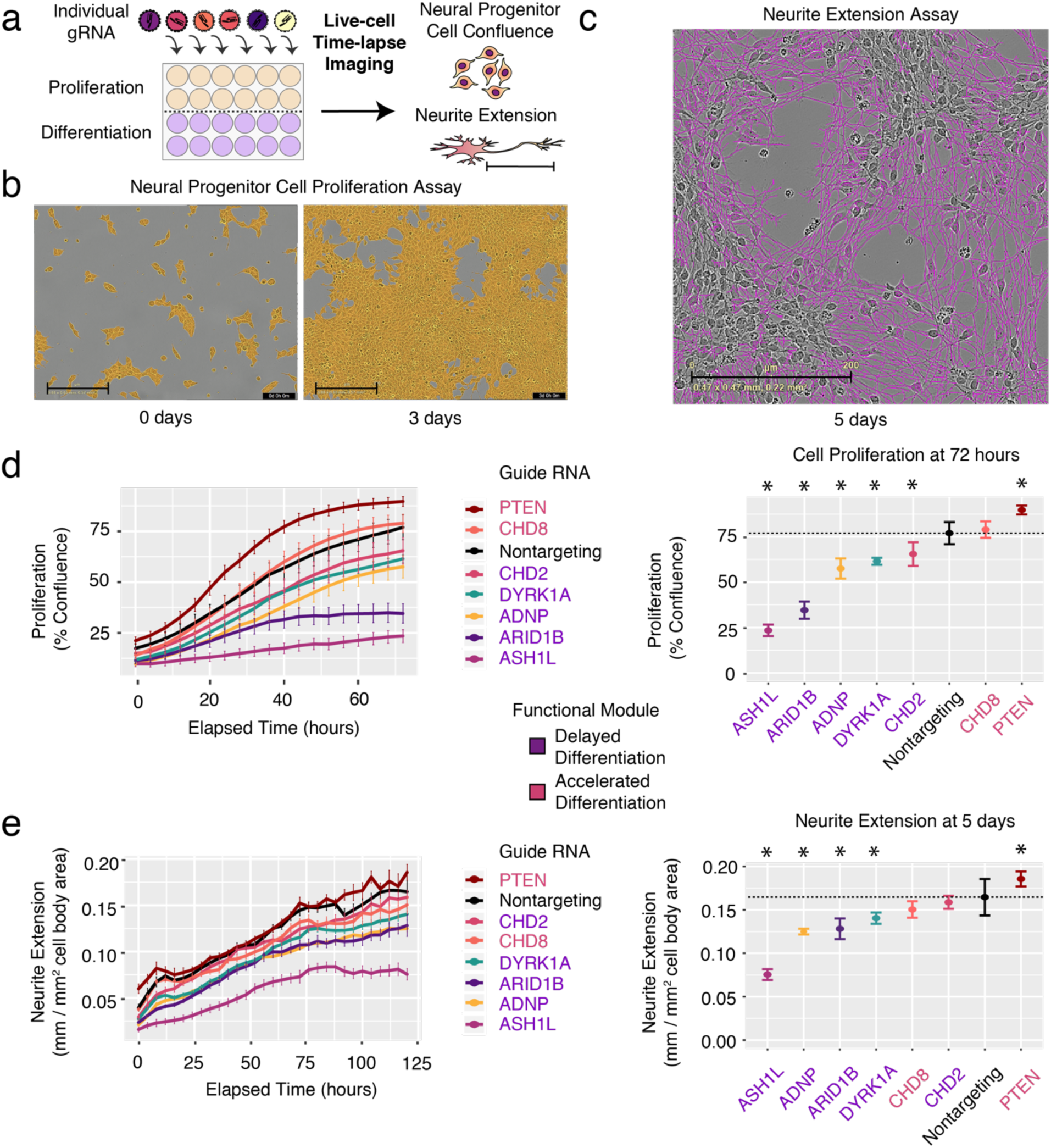
Live-cell imaging after repression of individual ASD genes confirms defects in cellular proliferation and neurite extension. **A**) Schematic overview of arrayed guide RNA (gRNA) screening. Cells infected with a single gRNA are assayed by time-course imaging for confluence and neurite extension. **B**) Representative images of neural progenitor cell proliferation assay at the start and end-points (day 0 and day 3). Neural progenitor cell proliferation is measured by creating a cell mask (orange) and computing the area of confluence at each time point. **C**) Representative image of neurite extension assay at 5 days post-differentiation. Neurite extension is measured with the NeuroTrack assay in the IncuCyte software. Neurite masks are shown (purple). Neurite extension lengths are normalized by cell cluster area to account for any differences in cell number. Scale bars = 200 µm. **d**) Time-lapse imaging of cellular proliferation (left), assessed by the percentage of confluence, reveals significant decreases or increases (right, * p < 0.01, t-test, dotted horizontal line indicates average in control cells). **E**) Time-lapse imaging of neurite extension (left) and quantification (right, * p < 0.01, t-test, dotted horizontal line indicates average in control cells). All values in (**d**) and (**e**) represent mean ± SEM. Cells with each individual gRNA were plated in duplicate or triplicate wells for each experiment. Images were captured from 9 fields per well at each timepoint. Experiments were repeated 2-3 times for all gRNAs.

Remarkably, in these live-cell imaging experiments, we observed decreased proliferation after repression of each of the 5 proposed ‘Delayed Differentiation’ genes (Fig 5d). In contrast, *PTEN* repression caused a major increase in proliferation, consistent with our prediction and with its known function as an inhibitor of neural stem cell proliferation^46^. For the neurite extension assay, most of our predictions were also confirmed (Fig 5e), as repression of 4 of the 5 ‘Delayed Differentiation’ genes, *ASH1L, ADNP, ARID1B*, and *DYRK1A*, caused modest to severe reductions in neurite outgrowth. In agreement with our transcriptome-based prediction, *PTEN* repression increased neurite extension in this assay, a result consistent with the earlier observation that *PTEN* enhances the length of regenerating axons *in vivo* ^47^. Together, proliferation and neurite extension assays confirmed the consequences of ASD-gene repression predicted by scRNA-seq analyses, demonstrating the utility of our approach for high-throughput functional elucidation of neurodevelopmental disease-associated genes. Live-cell imaging further supports the functional convergence of some ASD genes acting at an early stage to delay neuron differentiation and decrease proliferation.

### CRISPR repression in iPSC neural progenitor cells confirms functional gene modules and transcriptional convergence at cell-cycle dysregulation

To validate the early transcriptional convergence and effects on cellular proliferation of the ASD gene modules in an orthogonal cellular model system, we performed CRISPR repression of individual genes in human iPSC-derived neural progenitor cells. First, we confirmed that dCas9-KRAB repression was efficient in these cells for a subset of genes using qPCR (Figure 6a). Next, we performed RNA-seq after knockdown of seven individual ASD genes and a non-targeting control (Methods). RNA-seq confirmed efficient knockdown for 5/7 target genes (Supplementary Figure 7A). Clustering transcriptomes using principal component analysis closely reproduced the gene modules discovered in LUHMES, namely the clustering of four members of the ‘Delayed Differentiation’ gene set (*ADNP, ARID1B, ASH1L, and DYRK1A*) and the clustering of ‘Accelerated Differentiation’ genes *CHD8* and *PTEN* (Fig 6b). The same clusters were also observed by unsupervised hierarchical clustering of transcriptomes using highly variable genes (Supplementary Figure 7B). As in LUHMES, the ‘Delayed Differentiation’ genes strongly converged at the level of transcriptional regulation (Fig 6c), affecting genes enriched for roles in chromatin remodeling, Wnt signaling and cell-cycle regulation (Fig 6d). In these cells, the two ‘Accelerated Differentiation’ genes had strongly overlapping transcriptional consequences on both down- and up-regulated genes (Sup Fig 7C) leading to misregulation of cell-cycle genes and increased cell division pathways (Sup Fig 7D). This transcriptional convergence could be explained by the observation that *CHD8* repression also decreased the expression of *PTEN* (Figure 6A), implying that these genes are in the same pathway. A proliferation assay in these cells after individual gene repression functionally confirmed that *ASH1L* and *CHD2* decreased proliferation and *CHD8* repression enhanced it (Fig 6e). These results broadly confirm both the membership and functional interpretation of ASD-genes modules in a second human neural progenitor system, further validating the LUHMES as a relevant model for first pass high-throughput functional genomics screening.

**Figure 6:**
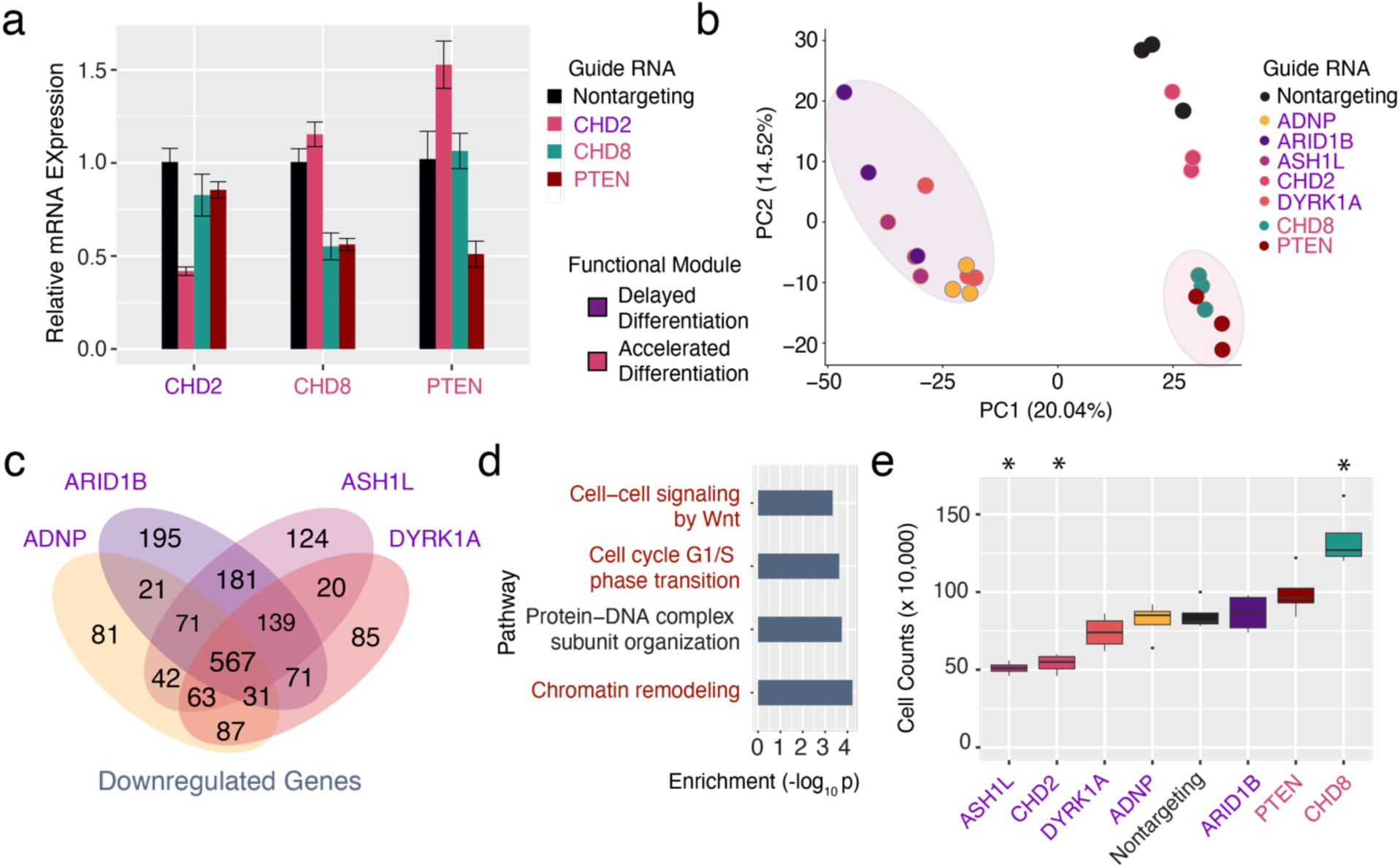
CRISPR repression in iPSC neural progenitor cells confirms modules of ASD genes and transcriptional convergence at cell-cycle dysregulation. **a**) Efficient dCas9-KRAB repression of individual target genes using the designated gRNAs in iPSC-NPCs. N = 3 biological replicates for all qPCR experiments. Values represent mean ± SEM. **b)** Clustering of RNA-seq profiles by principal component analysis reveals clustering of ‘Delayed’- and ‘Accelerated Differentiation’ module ASD genes **c)** ‘Delayed Differentiation’ module gene repression elicits strongly overlapping transcriptional consequences. **d)** Gene Ontology analysis of downregulated genes shows enrichment for chromatin remodeling and cell-cycle genes. **e)** Cellular proliferation measured by cell number after individual gene repression reveals significant decreases or increases (* indicates *p* < 0.01, n = 4).

### Functionally convergent ASD gene modules predict shared clinical phenotypes

Linking genotype to phenotype is the ultimate goal of functional genomics. To this end, we sought to determine if the functional convergence of ASD genes observed in our cellular model could predict a convergence of clinical phenotypes for these genes. To do so, we first integrated the results of our pseudotime analysis, transcriptional clustering, and functional profiling by hierarchical clustering (Figure 7a). As expected, this integrated model clearly separated the ‘Delayed Differentiation’ module genes from those in the ‘Accelerated Differentiation’ module. Next, we performed hierarchical clustering on the prevalence of clinical phenotypes from one study on individuals with dominant loss-of-function mutations in these genes^48^. Strikingly, clustering by clinical phenotypes fully recapitulated our proposed convergent modules and supports mechanistic links of convergent pathways to shared clinical outcomes (Fig 7b). For example, individuals with mutations in the ‘Delayed Differentiation’ module genes were highly likely to have intellectual disability, consistent with increased severity owing to early neurodevelopmental dysregulation. Comparing across the two clusters, these individuals have a higher incidence of microcephaly which is mechanistically consistent with the neural progenitor cell proliferation defects we observed. Likewise, individuals with *PTEN* and *CHD8* mutations have a comparatively reduced prevalence of intellectual disability but a high prevalence of macrocephaly, consistent with the observed functional convergence of these genes on promoting neuronal differentiation, proliferation, and neurite outgrowth. Recent studies have confirmed that disruptive mutations in *ADNP*, *ARID1B*, *CHD2* and *DYRK1A* are associated with a higher prevalence of severe neurodevelopmental delay. Conversely, *CHD8* and *PTEN* mutations are associated with ASD without neurodevelopmental delay^6^.

**Figure 7:**
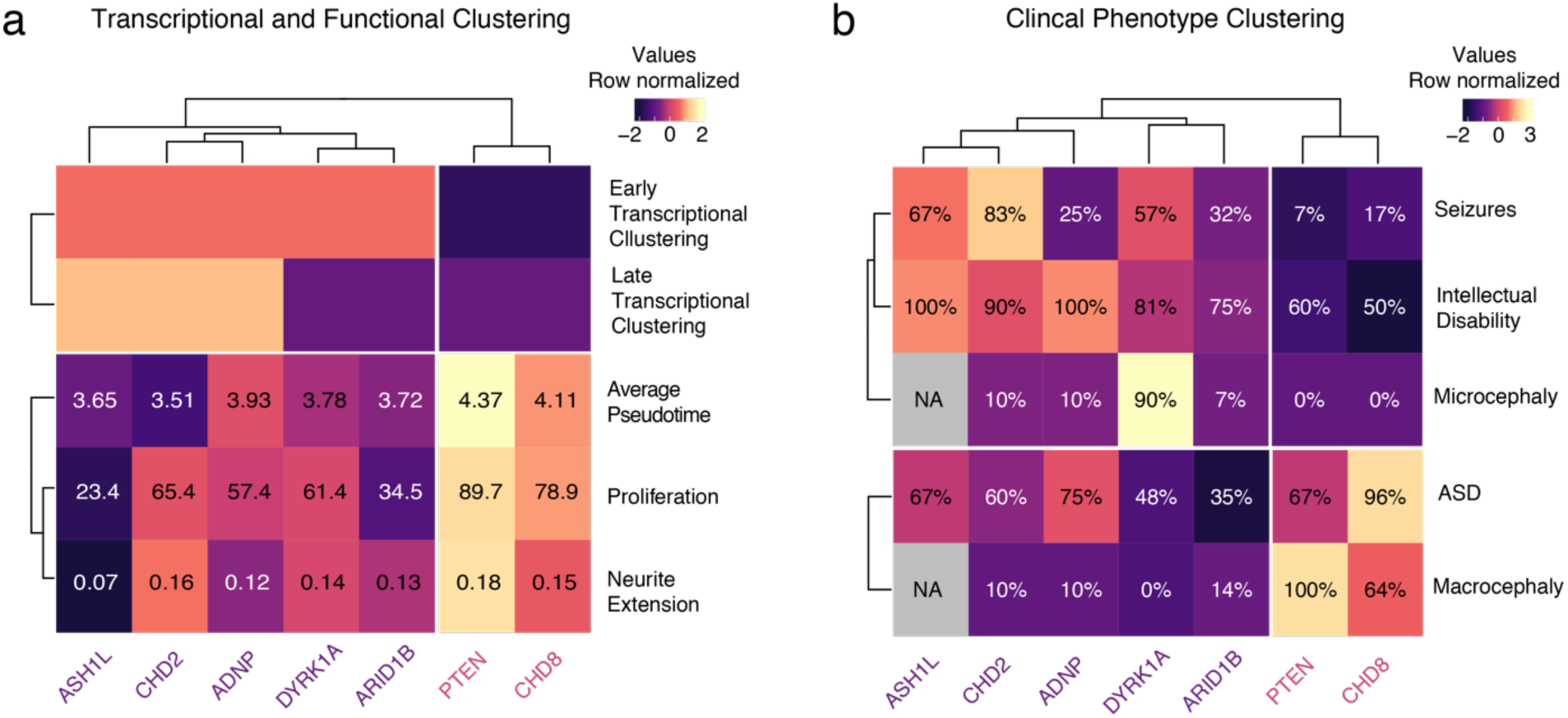
Experimental and clinical convergence of functional ASD gene modules. **A**) Integrating transcriptional and functional assays reveals and refines two functionally convergent modules of ASD genes. **b**) Clinical phenotype data reveals the same two modules of ASD genes. **%:** Prevalence of phenotype (percentage) in individuals from ^48^

## Discussion

The genetic and phenotypic heterogeneity of neurodevelopmental disorders challenge the understanding and treatment of these conditions. Over 1100 genes have been discovered to cause neurodevelopmental disorders when mutated and this number will continue to increase. As a result, it is imperative to develop better methods to cost-effectively unravel the functional contributions of these genes to both normal development and disease. Furthermore, the identification of convergent pathogenic mechanisms across diverse causative genes would facilitate the development of therapeutic interventions, but this requires a rapid, scalable, and disease-relevant model system in which tens to hundreds of genes can be modulated in parallel and their effects measured in a robust manner. Here we have taken an important step toward establishing such a system by coupling pooled dCas9-based transcriptional repression to single-cell RNA-sequencing in a simple and highly tractable human model of neuron differentiation. Using this approach, we perturbed the expression of 13 diverse autism-associated genes and uncovered unique and overlapping consequences on transcriptional networks and pathways. This led to specific predictions about functional roles of these genes in growth and neurite extension which we then validated through imaging. In addition to demonstrating the utility of a high-throughput functional genomics approach to dissect gene function, our results suggest that many ASD genes might act on common pathways, namely by modifying neural progenitor cell proliferation and cell-cycle.

By integrating pseudotime analysis, transcriptional clustering, and the cellular phenotyping of individual neurodevelopmental genes, we uncovered two consistent modules of genes with opposing functionality in altering the course of neuron differentiation. We identified a ‘Delayed Differentiation’ module comprised of 5 ASD-genes and predicted that the knockdown of these genes would decrease neural progenitor cell proliferation. We further predicted, for a subset of this ‘Delayed Differentiation’ module, that individual gene knockdown would decrease neuron projection. Finally, we also predicted that *PTEN* repression would increase both proliferation and neuron projection. Neural progenitor cell proliferation, the decision to exit the cell-cycle, and neuron differentiation are complex and interrelated processes^49^. Increased proliferation could either drive an expansion of a progenitor pool or increase neurogenic cell divisions leading to precocious differentiation. A parsimonious explanation of our data in LUHMES is that the progression of neuron differentiation is preceded by a few requisite neurogenic cell divisions. Delaying these divisions by disrupting cell cycle, or enhancing them by increasing proliferation rate, will consequently delay or accelerate neuronal differentiation progression respectively. We confirmed these predictions by performing live-cell imaging of cell proliferation and neurite extension after gene knockdown, providing experimental functional validation for these ASD genes. Convergence of ASD genes at the regulation of cell-cycle genes was supported in an orthogonal model using human iPSC-derived NPCs. Heterochronicity of neurodevelopmental gene expression networks and consequent dysregulation of neuron differentiation is a plausible mechanism underlying ASD pathology, and has been observed in other cellular models of ASD^13, 50^.

We have demonstrated that LUHMES cells enable the rapid and robust generation of post-mitotic human neurons with transcriptional profiles that correspond closely to early human cortical development, a critical period of neurodevelopment that has been implicated in the etiology of ASD and other neurodevelopmental disorders. Confirmation of our results in human iPSC-derived NPCs further validates the utility of LUHMES for discovering potential mechanisms of a subset of neurodevelopmental disease-associated genes. Because transcriptional regulators are particularly enriched for high early-fetal expression^6^, LUHMES may be especially suited to studying this class of genes. However, for some other classes of neurodevelopmental genes, LUHMES cells may not be the most well-suited model system. For example, synaptic genes are more postnatally expressed across a broad range of neuronal types. iPSC-derived induced excitatory and inhibitory neurons, co-cultures of these cells, or 3D-organoids are likely to be more suitable than LUHMES for studying these genes and their roles in establishing synaptic phenotypes. These alternative human neurodevelopmental models may also be better suited for investigating genes and pathways involved in cell-fate specification, neuronal migration, and neuronal activity. However, the experimental complexity and heterogeneity of these models compared to the rapid and reproducible differentiation of LUHMES make them less suited for high-throughput analyses of sets of genes that are likely to be involved in neuronal differentiation or maturation.

Compared to traditional single gene disease modeling experiments in mice or human cells (*e.g.* refs. ^51–55^), our approach increases the number of genes that can be assayed in parallel while also overcoming many of the primary sources of variation in such models. Critically, this enables direct comparison of results across genes to discover convergent mechanisms. Current single-cell technology enables pathway-level inferences of transcriptional dysregulation and prioritization of candidate genes for further functional validation in a rapid and cost-effective manner for tens of disease implicated genes. However, noise and sparsity in single cell RNA-seq data limit its power to detect differentially expressed genes. Quantitative measurements of knockdown efficiency of individual gRNAs within single cells are difficult, and individually validating gRNAs is incompatible with the goal of high-throughput screening. Our qPCR measurements of gRNA efficiencies showed a range of 40-90% knockdown for all tested target genes, which may better represent haploinsufficiency than a total knock-out approach. Exquisite dosage sensitivity of certain genes raises the possibility of variable cellular phenotypes depending on the degree of knockdown. Including more gRNAs per gene, and enough cells to analyze effects on a per-gRNA instead of per-gene basis, would resolve these potential confounds. Improvements in the sensitivity and throughput of scRNA-seq, as well as declining costs, will enhance its utility in future experiments.

A major strength of our approach is its extensibility to different model systems for the rapid functional profiling of diverse gene sets, allowing for prioritization of candidate genes for low-throughput cellular phenotyping by imaging. Although we only measured proliferation and neurite extension with live-cell imaging, the integration of these data with pooled transcriptomes revealed consistent gene modules that were also convergent at the level of clinical phenotypes in individuals with mutations in these genes^48^, illustrating the potential of this approach for linking molecular pathways to clinical phenotypes. Furthermore, while not implemented in this study, this approach is amenable to screening of chemical libraries to discover effective pharmaceutical interventions for any observed defects, and to determine whether convergent genetic modules will respond to common treatments. Pooled, rather than arrayed, optical phenotyping approaches would further accelerate these efforts^56^. Such high-content imaging and screening in future experiments will enable detailed characterization of perturbation-induced neuronal phenotypes and the discovery of convergent molecular endophenotypes of disease pathogenesis.

## Materials and Methods

### Cell culture

HEK293T cells were maintained in Dulbecco’s Modified Eagle Media (DMEM) supplemented with 10% fetal bovine serum and 1% penicillin-streptomycin, and passaged every 3-4 days after enzymatic dissociation using trypsin. LUHMES cells (ATCC cat# CRL-2927) were cultured according to established protocols with minor modifications ^17^. Proliferating LUHMES cells were maintained in DMEM/F12 media supplemented with 1% N2 and 40 ng/mL basic fibroblast growth factor (bFGF) and 1% penicillin-streptomycin. Cells were grown in T25 flasks coated with poly-ornithine and fibronectin. For differentiation, bFGF was withdrawn from the media and tetracycline was added (1 µg /mL) to repress the v-Myc transgene. For time-course RNA-sequencing experiments, neurotrophic factors (cAMP and GDNF) were added to the differentiation media. These factors were withheld from repression experiments to increase the sensitivity to detect perturbations. For time-course differentiation, two replicates of LUHMES cells were differentiated for each time point and total mRNA was purified. RNA was sent for sequencing at the Genome Technology Access Center (GTAC) at Washington University School of Medicine. Polyclonal dCas9-KRAB-blast expressing LUHMES were generated by infecting cells with lentivirus and selecting using blasticidin (10 µg/mL). Lenti-dCas9-KRAB-blast was a gift from Gary Hon (Addgene #89567)^57^. dCas9-KRAB LUHMES were maintained in blasticidin-containing media to prevent transgene silencing.

Human iPSC-derived neural progenitor cells (XCL4) were acquired from STEMCELL technologies (cat #70902) and grown in Neural Progenitor Medium 2. While now discontinued by STEMCELL technologies, these reagents are available from XCellScience. Tetracycline inducible dCas9-KRAB NPCs were generated after neomycin selection (200 µg/mL). pHAGE TRE dCas9-KRAB was a gift from Rene Maehr and Scot Wolfe (Addgene #50917)^58^. Cells were plated at a density of 200,000 cells per well in a 12-well plate on Matrigel and infected in triplicate with individual guide RNAs targeting ASD genes. Next day, media containing doxycycline (2µg/mL) and puromycin (1 µg/mL) was added to induce dCas9-KRAB and select for guide-containing cells. Cells were passaged 4 days after selection. mRNA was collected 4 days after replating (8 days of repression), and 3’ RNA-seq libraries were prepared by BRB-seq^59^. For proliferation assays, TRE dCas9-KRAB XCL4 cells were infected in quadruplicate with gRNAs targeting 7 ASD-genes and 1 non-targeting gRNA. After puromycin selection and doxycycline induction, cells were plated at equal cell numbers, grown for 8 days, and total cells were counted using a hemocytometer. Cell counts were compared to cells infected with a nontargeting gRNA

### gRNA Cloning

For each target gene, we selected three gRNAs optimized for repression from the Dolcetto library^36^. These gRNAs direct dead Cas9 (dCas9) to a window 25-75 nucleotides downstream of the gene’s transcription start site. gRNAs were screened for sequence features predicting high activity and no off-target effects. gRNAs were cloned into a CRISPR-repression optimized vector to enable pooled lentiviral preparation without guide-barcode swapping^35–37^. Lentiviral gRNA expression vectors were created by annealing two complementary oligonucleotides encoding gRNAs at 100 µM (IDT DNA) with sticky-ends and ligating annealed products into BsmB1 digested CROP-seq-opti vector using Golden Gate assembly. CROP-seq-opti was a gift from Jay Shendure (Addgene #106280)^35^. Each Golden Gate assembly reaction contained 6.5 µL water, 1 µL 1:10 diluted annealed oligos, 1 µl T4 ligase, 1 µl T4 ligase buffer, and 0.5 µL BsmB1. Reactions were incubated at 16 °C for 10 minutes (ligation) and 55 °C for 10 minutes (restriction) for 4 cycles. 1 µL of Golden Gate mixture was transformed into 30 µL Stellar Competent cells and plated onto ampicillin-containing agar plates. Individual colonies were miniprepped after colony PCR and all constructs were verified by Sanger sequencing (Genewiz).

### Lentivirus production of individual gRNAs and pooled gRNA libraries

Lentivirus was produced according to established protocols. In brief, HEK293T cells were seeded at a density of one million cells per well of a 6 well plate and transfected with 2 micrograms of DNA comprising 1 µg gRNA-transfer plasmid, 750 ng psPAX2 and 250 ng pMD2.G. psPAX2 and pMD2.G were gifts from Didier Trono (Addgene #12259 & #12260). Cells were transfected using the PEI method (Polysciences). Media was changed 12 hours after transfection and viral-containing supernatant was collected 24 and 48 hours later. Lentivirus was concentrated using LentiX reagent and resuspended in 50 µL aliquots for each mL of original supernatant (20X concentration). Lentivirus was titered on LUHMES cells by infecting cells with serial dilutions of virus, followed by antibiotic selection (puromycin for gRNAs, 1 µg / mL). For pooled gRNA libraries, equal amounts of DNA for each gRNA were mixed prior to transfection.

### Lentiviral transduction of gRNAs

For individual or pooled gRNAs, LUHMES cells were infected with serial dilutions of virus. Virus-containing media was removed after 4-6 hours of transduction. Antibiotic selection with puromycin (1 µg/mL) was applied 24 hours after infection. Wells in which no more than 25% of cells survived, corresponding to multiplicity of infection < 0.3, were used for experiments. Cells were expanded for 4 days before plating for differentiation. After re-plating, cells were differentiated for 6 to 8 days in the presence of tetracycline to allow efficient repression and differentiation. Puromycin and blasticidin were maintained for the duration of all experiments to ensure gRNA and dCas9-KRAB expression in all cells.

### Quantitative real-time PCR

RNA was purified from 6-8 day differentiated LUHMES using Trizol. 1 µg of RNA was reverse-transcribed into cDNA using qScript cDNA Supermix (Quantabio). Quantitative real-time PCR (qRT-PCR) was performed on an ABI 7900HT using Sybr Green SuperMix (Quantabio). Relative expression levels were determined using the comparative threshold (ΔΔCT) method^60^. Beta-actin (ACTB) mRNA levels were used as a normalization control. Sequences for qRT-PCR primers are:

ADNP qPCR fw: CATGGGAGGATGTAGGACTGT

ADNP qPCR rv: ATGGACATTGCGGAAATGACT

CTNND2 qPCR fw: AGGTCCCCGTCCATTGATAG

CTNND2 qPCR rv: ACTGGTGCTGCAACATCTGAA

PTEN qPCR fw: TTTGAAGACCATAACCCACCAC

PTEN qPCR rv: ATTACACCAGTTCGTCCCTTTC

DYRK1A qPCR fw: AAGAAGCGAAGACACCAACAG

DYRK1A qPCR rv: TTTCGTAACGATCCATCCACTTT

CHD8 qPCR fw: CTGCACAGTCACCTCGAGAA

CHD8 qPCR rv: TGGTTCTTGCACTGGTTCAG

HDAC5 qPCR fw: GTACCCAGTCCTCCCCTGC

HDAC5 qPCR rv: GCACATGCACTGGTGCTTTA

ACTB qPCR fw: CATGTACGTTGCTATCCAGGC

ACTB-qPCR rv: CTCCTTAATGTCACGCACGAT

### Single cell transcriptome capture

12,000 cells were loaded per lane of a 10X Chromium device using 10X V2 Single Cell 3ʹ Solution reagents (10X Genomics, Inc). Two biological replicates of pooled single-cell experiments were performed independently. Each replicate was loaded across 1 or 2 lanes of a 10X Single Cell A Chip V2. Single cell libraries were prepared following the Single Cell 3’ Reagent Kits v2 User Guide (Rev B). Single cell cDNA libraries were amplified for 12 initial cycles after reverse transcription. A fraction of the pre-fragmented cDNA libraries was reserved for gRNA-specific enrichment PCR.

### gRNA-transcript enrichment PCR

Three gRNA-specific enrichment PCR replicates were performed for each single cell library. Each reaction used 1 µL of the single-cell libraries as a template to amplify captured gRNA sequences. A single-step PCR reaction was used to amplify gRNA from total captured cDNA libraries using custom primers:

P5-index-Seq1-fw: AATGATACGGCGACCACCGAGATCTACACAGGACAACACTCTTTCCCTACACGACGCTCTTCCG ATCT

P7-Index-Seq2-10x-sgRNA:

CAAGCAGAAGACGGCATACGAGATCGGGCAACGGTGACTGGAGTTCAGACGTGTGCTCTTCCG ATCTGTGGAAAGGACGAAACA*C*C*G

* denotes phosphorothioate modification to reduce mispriming due to proof-reading polymerase

### gRNA Depletion Analysis

Almost all of the gRNAs in our lentiviral pool (43/47) were well-represented in perturbed cells at similar frequencies, yet 4 gRNAs (targeting *ASH1L*, *POGZ gRNAs 1 and 2, and SETD5*) were significantly depleted (chi-squared test, *p* < 0.01, Supplementary Figure 8A). We hypothesized that their depletion was the result of fitness defects caused by the repression of these genes^61, 62^. To test this, we infected neural progenitor cells individually with three of the depleted gRNAs and monitored cell proliferation using live-imaging. Compared to a non-targeting gRNA, all of the depleted gRNAs caused a significant reduction in cellular proliferation, explaining why few cells with these gRNAs were detected in our pooled experiment (Supp Fig 8B).

### Bioinformatic Analyses

Sequencing data corresponding to single-cell transcriptomes were processed using the 10X software package Cell Ranger (v 2.1.0). We used this software to map reads to hg38 using STAR (v2.5.1b). The output filtered gene expression matrices were imported into R (v 3.5.1) for further analysis. Most analyses were performed using the Seurat (v3.0) and Monocle (v2.10.0) packages. Individual cells were sequenced to an average depth of 50,208 ± 7,310 mapped reads per cell (2,145 ± 448 genes detected, 6,625 ± 1766 unique molecular identifiers (UMIs)). Quality control was performed in Seurat by computing the number of transcripts per cell and the percentage of mitochondrial gene expression. Cells with more than 500 but fewer than 7500 detected genes, and less than 8% mitochondrial gene expression were retained. gRNAs were detected by next-generation sequencing of the custom gRNA-enrichment PCR. Look-up tables of gRNA and cell barcodes were generated using custom Python scripts with a detection threshold of 20 UMIs per gRNA-cell barcode combination. Cell barcodes from filtered high-quality cells were matched against this table and only cells with a single gRNA were retained for analysis. We estimated gRNA knock-down efficiency separately on independent replicates using the mimosca.run_model() command in the MIMOSCA toolkit. Differences in total UMIs, experimental batch, and mitochondrial percentage were accounted for during data normalization. Normalized filtered data were used for the remaining analyses. For each perturbation, we grouped all cells with any of the three gRNAs targeting the same gene, as we have demonstrated that all three gRNAs typically have strong on-target activity (Sup Fig 2D).

### Clustering Analysis

Dimensionality reduction was performed by running principal component analysis (PCA) and clustering cells by the first 6 PCAs using UMAP^63^. To assess global variation in transcriptional states, we visualized all single cell transcriptomes on the UMAP. This revealed that over 99% of all cells formed a single cluster of post-mitotic neurons as defined by the absence of proliferative marker expression (Sup Fig 3A-B). This is consistent with our experimental design capturing a single timepoint (day 7) in a rapid isogenic model of neuronal differentiation. Within the major cluster, however, the expression of markers of neuronal differentiation showed variable patterns across the UMAP (Sup Fig 3C-F). Moreover, the most variably expressed genes in single-cell transcriptomes were enriched for functional roles in neurogenesis and axon projection, suggesting heterogeneity of neuronal differentiation at the single-cell level.

### Pseudotime Analysis

We projected cells in pseudotime in Monocle by ordering cells by highly variable genes. Dimensionality reduction was performed using the “DDRTree” method. Trajectories based on different sets of highly variable genes were qualitatively similar, showing a single trajectory with only minor branching. The pseudotime trajectory is comprised of individual line segments called pseudotime ‘States’. To ensure high correlation between pseudotime and neuron differentiation status, we computed the state-specific genes in the bulk RNA-seq dataset for each day of differentiation and used these genes for pseudotime ordering. We then transferred pseudotime state labels into Seurat to discover marker genes for each pseudotime state. Transferring pseudotime labels onto the UMAP plot showed distinct banding patterns representing subtle transcriptional state differences within the main cluster (Supplementary Figure 9A). Re-clustering the earliest and latest pseudotime labeled cells revealed two completely distinct cell states expressing either differentiation (*NEUROD1*) or maturation (*STMN2*) markers^64, 65^ (Supp Fig 9C-E). This confirms that pseudotime is more sensitive to detect biologically relevant transcriptional patterns than UMAP clustering in our dataset. We tested for altered pseudotime state membership proportions for each gRNA using chi-squared test. We computed the distribution of pseudotime state scores for each gRNA, and compared their averages using t-tests.

### Transition Mapping of LUHMES Differentiation

Transition mapping allows the comparison of *in vitro* neuron differentiation to *in vivo* development by computing the overlap of differentially expressed genes at selected time points across datasets^24^. We compared the *in vitro* LUHMES differentiation timepoints day 0 to day 8 to transcriptional changes across brain regions and developmental timepoints in the BrainSpan Atlas of the Developing Human Brain. LUHMES differentiation had the strongest overlap with transcriptional changes occurring in the cortex of post-conception week 8-10 embryos and week 10-13 embryos.

### Differential Gene Expression Analysis

Differential gene expression testing was performed using the FindMarkerGenes function in Seurat using the Wilcoxon Rank Sum test and a relaxed log2FC threshold of 0.1 to increase the number of differentially expressed genes. This cutoff was calibrated against a ‘gold standard’ dataset comparing single-cell and bulk RNA-seq data to identify differentially expressed genes^66^. To find marker genes of pseudotime state clusters, only positive markers were returned.

Pseudotime state by was binarized with states 1-3 labeled as ‘early’ and states 4-6 as ‘late’. We next created another label combining the targeted gene with binary pseudotime state (e.g. CHD8_early). Averaged profiles were re-computed for each group based on these new labels and differentially expressed genes were also re-calculated. Principal component analysis and hierarchical clustering were performed on these samples. As expected, unsupervised clustering of the stratified profiles by principal component analysis perfectly discriminated between ‘early’ and ‘late’ samples (Sup Fig 5C). This analysis showed that the first principal component corresponds to pseudotime status and explains almost 20% of the total variance in the dataset. Gene Ontology and pathway enrichment analyses were performed using WebGestalt^67^.

### Live-cell Imaging

Cells were imaged using an IncuCyte S3 live imaging system (Essen BioScience) For each experiment, dCas9-KRAB LUHMES were infected in duplicate or triplicate with individual gRNAs and were plated in duplicate or triplicate in wells of a 24 well plate in either self-renewing or differentiation conditions. 9 fields per well were imaged every 4 hours for either 3 or 5 days for proliferation or differentiation respectively. These experiments were repeated 2-3 times for each individual gRNA. Images were analyzed using the IncuCyte Software. Specifically, we performed the Proliferation Analysis and NeuroTrack neurite tracing analyses with default parameters. Cell bodies and neurites were detected from phase contrast images. Representative images are shown in Supplementary Figure 4.

### Data Availability

Sequencing data generated in this study are available through GEO (GSE142078).

## Supporting information

Lalli et al Supplementary Figures

## Acknowledgments

This work was supported by:

T32HL125241-05 (NHLBI)

U54HD087011-04 (NICHD)

R21NS087230-01A1 (NINDS) (RM & JM)

RF1MH117070 and U01MH109133 (NIMH)(RM and JD)

SFARI Explorer #: 500661 (RM & JD)

T32CA009547-33 (NCI) (DA)

GTAC is partially supported by National Institutes of Health (NIH) grants P30 CA91842 and UL1 TR000448

## Author contributions

M.A.L. performed experiments, analyzed data, designed the study, and wrote the manuscript. D.R.A. performed experiments and wrote the manuscript. J.D.D., J.M., and R.D.M analyzed data, designed the study, and wrote the manuscript.

## References

1. Amberger, J. S., Bocchini, C. A., Scott, A. F. & Hamosh, A. OMIM.org: leveraging knowledge across phenotype–gene relationships. Nucleic Acids Res. 47, D1038–D1043 (2019).

2. Wright, C. F. et al. Genetic diagnosis of developmental disorders in the DDD study: a scalable analysis of genome-wide research data. The Lancet 385, 1305–1314 (2015).

3. O’Roak, B. J. et al. Sporadic autism exomes reveal a highly interconnected protein network of *de novo* mutations. Nature 485, 246–250 (2012).

4. De Rubeis, S. et al. Synaptic, transcriptional and chromatin genes disrupted in autism. Nature 515, 209–215 (2014).

5. Iossifov, I. et al. The contribution of de novo coding mutations to autism spectrum disorder. Nature 515, 216–221 (2014).

6. Satterstrom, F. K. et al. Large-Scale Exome Sequencing Study Implicates Both Developmental and Functional Changes in the Neurobiology of Autism. Cell 0, (2020).

7. Kilpinen, H. et al. Common genetic variation drives molecular heterogeneity in human iPSCs. Nature 546, 370–375 (2017).

8. Zhao, X. & Bhattacharyya, A. Human Models Are Needed for Studying Human Neurodevelopmental Disorders. Am. J. Hum. Genet. 103, 829–857 (2018).

9. Qi, L. S. et al. Repurposing CRISPR as an RNA-Guided Platform for Sequence-Specific Control of Gene Expression. Cell 152, 1173–1183 (2013).

10. Adamson, B. et al. A Multiplexed Single-Cell CRISPR Screening Platform Enables Systematic Dissection of the Unfolded Protein Response. Cell 167, 1867–1882.e21 (2016).

11. Dixit, A. et al. Perturb-Seq: Dissecting Molecular Circuits with Scalable Single-Cell RNA Profiling of Pooled Genetic Screens. Cell 167, 1853–1866.e17 (2016).

12. Datlinger, P. et al. Pooled CRISPR screening with single-cell transcriptome readout. Nat. Methods 14, 297–301 (2017).

13. Schafer, S. T. et al. Pathological priming causes developmental gene network heterochronicity in autistic subject-derived neurons. Nat. Neurosci. 1 (2019) doi:10.1038/s41593-018-0295-x.

14. Willsey, A. J. et al. The Psychiatric Cell Map Initiative: A Convergent Systems Biological Approach to Illuminating Key Molecular Pathways in Neuropsychiatric Disorders. Cell 174, 505–520 (2018).

15. Ho, S.-M. et al. Evaluating Synthetic Activation and Repression of Neuropsychiatric-Related Genes in hiPSC-Derived NPCs, Neurons, and Astrocytes. Stem Cell Rep. 9, 615–628 (2017).

16. Hoffman, G. E. et al. Transcriptional signatures of schizophrenia in hiPSC-derived NPCs and neurons are concordant with post-mortem adult brains. Nat. Commun. 8, 2225 (2017).

17. Scholz, D. et al. Rapid, complete and large-scale generation of post-mitotic neurons from the human LUHMES cell line. J. Neurochem. 119, 957–971 (2011).

18. Höllerhage, M. et al. Protective efficacy of phosphodiesterase-1 inhibition against alpha-synuclein toxicity revealed by compound screening in LUHMES cells. Sci. Rep. 7, 11469 (2017).

19. Tong, Z.-B. et al. Characterization of three human cell line models for high-throughput neuronal cytotoxicity screening. J. Appl. Toxicol. 37, 167–180 (2017).

20. Pierce, S. E., Tyson, T., Booms, A., Prahl, J. & Coetzee, G. A. Parkinson’s disease genetic risk in a midbrain neuronal cell line. Neurobiol. Dis. 114, 53–64 (2018).

21. Shah, R. R. et al. Efficient and versatile CRISPR engineering of human neurons in culture to model neurological disorders. Wellcome Open Res. 1, (2016).

22. Matelski, L., Morgan, R. K., Grodzki, A. C., Van de Water, J. & Lein, P. J. Effects of cytokines on nuclear factor-kappa B, cell viability, and synaptic connectivity in a human neuronal cell line. Mol. Psychiatry 1–13 (2020) doi:10.1038/s41380-020-0647-2.

23. Xu, X., Wells, A. B., O’Brien, D. R., Nehorai, A. & Dougherty, J. D. Cell Type-Specific Expression Analysis to Identify Putative Cellular Mechanisms for Neurogenetic Disorders. J. Neurosci. 34, 1420–1431 (2014).

24. Stein, J. L. et al. A Quantitative Framework to Evaluate Modeling of Cortical Development by Neural Stem Cells. Neuron 83, 69–86 (2014).

25. BrainSpan Atlas of the Developing Human Brain [Internet]. Funded by ARRA Awards 1RC2MH089921-01, 1RC2MH090047-01, and 1RC2MH089929-01. ©2011. Available from: http://brainspan.org/.

26. Li, M. et al. Integrative functional genomic analysis of human brain development and neuropsychiatric risks. Science 362, eaat7615 (2018).

27. Abrahams, B. S. et al. SFARI Gene 2.0: a community-driven knowledgebase for the autism spectrum disorders (ASDs). Mol. Autism 4, 36 (2013).

28. Amir, R. E. et al. Rett syndrome is caused by mutations in X-linked *MECP2*, encoding methyl-CpG-binding protein 2. Nat. Genet. 23, 185–188 (1999).

29. Medina, M., Marinescu, R. C., Overhauser, J. & Kosik, K. S. Hemizygosity of δ-Catenin (CTNND2) Is Associated with Severe Mental Retardation in Cri-du-Chat Syndrome. Genomics 63, 157–164 (2000).

30. Sun, G., Yu, R. T., Evans, R. M. & Shi, Y. Orphan nuclear receptor TLX recruits histone deacetylases to repress transcription and regulate neural stem cell proliferation. Proc. Natl. Acad. Sci. 104, 15282–15287 (2007).

31. Karczewski, K. J. et al. Variation across 141,456 human exomes and genomes reveals the spectrum of loss-of-function intolerance across human protein-coding genes. bioRxiv 531210 (2019) doi:10.1101/531210.

32. Lek, M. et al. Analysis of protein-coding genetic variation in 60,706 humans. Nature 536, 285–291 (2016).

33. Parikshak, N. N. et al. Integrative Functional Genomic Analyses Implicate Specific Molecular Pathways and Circuits in Autism. Cell 155, 1008–1021 (2013).

34. O’Leary, N. A. et al. Reference sequence (RefSeq) database at NCBI: current status, taxonomic expansion, and functional annotation. Nucleic Acids Res. 44, D733–D745 (2016).

35. Hill, A. J. et al. On the design of CRISPR-based single-cell molecular screens. Nat. Methods 15, 271–274 (2018).

36. Sanson, K. R. et al. Optimized libraries for CRISPR-Cas9 genetic screens with multiple modalities. Nat. Commun. 9, 5416 (2018).

37. Xie, S., Cooley, A., Armendariz, D., Zhou, P. & Hon, G. C. Frequent sgRNA-barcode recombination in single-cell perturbation assays. PLOS ONE 13, e0198635 (2018).

38. Zheng, G. X. Y. et al. Massively parallel digital transcriptional profiling of single cells. Nat. Commun. 8, 14049 (2017).

39. Satija, R., Farrell, J. A., Gennert, D., Schier, A. F. & Regev, A. Spatial reconstruction of single-cell gene expression data. Nat. Biotechnol. 33, 495–502 (2015).

40. Stuart, T. et al. Comprehensive Integration of Single-Cell Data. Cell 177, 1888–1902.e21 (2019).

41. Trapnell, C. et al. The dynamics and regulators of cell fate decisions are revealed by pseudotemporal ordering of single cells. Nat. Biotechnol. 32, 381–386 (2014).

42. McFaline-Figueroa, J. L. et al. A pooled single-cell genetic screen identifies regulatory checkpoints in the continuum of the epithelial-to-mesenchymal transition. Nat. Genet. 51, 1389–1398 (2019).

43. Duan, B. et al. Model-based understanding of single-cell CRISPR screening. Nat. Commun. 10, (2019).

44. Yang, L. et al. scMAGeCK links genotypes with multiple phenotypes in single-cell CRISPR screens. Genome Biol. 21, 19 (2020).

45. Vieth, B., Ziegenhain, C., Parekh, S., Enard, W. & Hellmann, I. powsimR: power analysis for bulk and single cell RNA-seq experiments. Bioinformatics 33, 3486–3488 (2017).

46. Groszer, M. et al. PTEN negatively regulates neural stem cell self-renewal by modulating G0-G1 cell cycle entry. Proc. Natl. Acad. Sci. 103, 111–116 (2006).

47. Park, K. K. et al. Promoting Axon Regeneration in the Adult CNS by Modulation of the PTEN/mTOR Pathway. Science 322, 963–966 (2008).

48. Stessman, H. A. F. et al. Targeted sequencing identifies 91 neurodevelopmental-disorder risk genes with autism and developmental-disability biases. Nat. Genet. advance online publication, (2017).

49. Hardwick, L. J. A. & Philpott, A. Nervous decision-making: to divide or differentiate. Trends Genet. 30, 254–261 (2014).

50. Marchetto, M. C. et al. Altered proliferation and networks in neural cells derived from idiopathic autistic individuals. Mol. Psychiatry (2016) doi:10.1038/mp.2016.95.

51. Cotney, J. et al. The autism-associated chromatin modifier CHD8 regulates other autism risk genes during human neurodevelopment. Nat. Commun. 6, ncomms7404 (2015).

52. Durak, O. et al. Chd8 mediates cortical neurogenesis via transcriptional regulation of cell cycle and Wnt signaling. Nat. Neurosci. 19, 1477–1488 (2016).

53. Katayama, S., Moriguchi, T., Ohtsu, N. & Kondo, T. A Powerful CRISPR/Cas9-Based Method for Targeted Transcriptional Activation. Angew. Chem. Int. Ed. 55, 6452–6456 (2016).

54. Bellmaine, S. F. et al. Inhibition of DYRK1A disrupts neural lineage specification in human pluripotent stem cells. eLife 6, e24502 (2017).

55. Jung, E.-M. et al. Arid1b haploinsufficiency disrupts cortical interneuron development and mouse behavior. Nat. Neurosci. 20, 1694 (2017).

56. Feldman, D. et al. Optical Pooled Screens in Human Cells. Cell 179, 787–799.e17 (2019).

57. Xie, S., Duan, J., Li, B., Zhou, P. & Hon, G. C. Multiplexed Engineering and Analysis of Combinatorial Enhancer Activity in Single Cells. Mol. Cell 66, 285–299.e5 (2017).

58. Kearns, N. A. et al. Cas9 effector-mediated regulation of transcription and differentiation in human pluripotent stem cells. Development 141, 219–223 (2014).

59. Alpern, D. et al. BRB-seq: ultra-affordable high-throughput transcriptomics enabled by bulk RNA barcoding and sequencing. Genome Biol. 20, 71 (2019).

60. Livak, K. J. & Schmittgen, T. D. Analysis of Relative Gene Expression Data Using Real-Time Quantitative PCR and the 2−ΔΔCT Method. Methods 25, 402–408 (2001).

61. Shalem, O. et al. Genome-Scale CRISPR-Cas9 Knockout Screening in Human Cells. Science 343, 84–87 (2014).

62. Wang, T., Wei, J. J., Sabatini, D. M. & Lander, E. S. Genetic Screens in Human Cells Using the CRISPR-Cas9 System. Science 343, 80–84 (2014).

63. Becht, E. et al. Dimensionality reduction for visualizing single-cell data using UMAP. Nat. Biotechnol. (2018) doi:10.1038/nbt.4314.

64. Polioudakis, D. et al. A Single-Cell Transcriptomic Atlas of Human Neocortical Development during Mid-gestation. Neuron 0, (2019).

65. Dennis, D. J., Han, S. & Schuurmans, C. bHLH transcription factors in neural development, disease, and reprogramming. Brain Res. 1705, 48–65 (2019).

66. Avey, D. et al. Single-Cell RNA-Seq Uncovers a Robust Transcriptional Response to Morphine by Glia. Cell Rep. 24, 3619–3629.e4 (2018).

67. Wang, J., Vasaikar, S., Shi, Z., Greer, M. & Zhang, B. WebGestalt 2017: a more comprehensive, powerful, flexible and interactive gene set enrichment analysis toolkit. Nucleic Acids Res. 45, W130–W137 (2017).

