## Supplementary material for "High-throughput single-cell functional elucidation of neurodevelopmental disease-associated genes reveals convergent mechanisms altering neuronal differentiation": Lalli et al Supplementary Figures

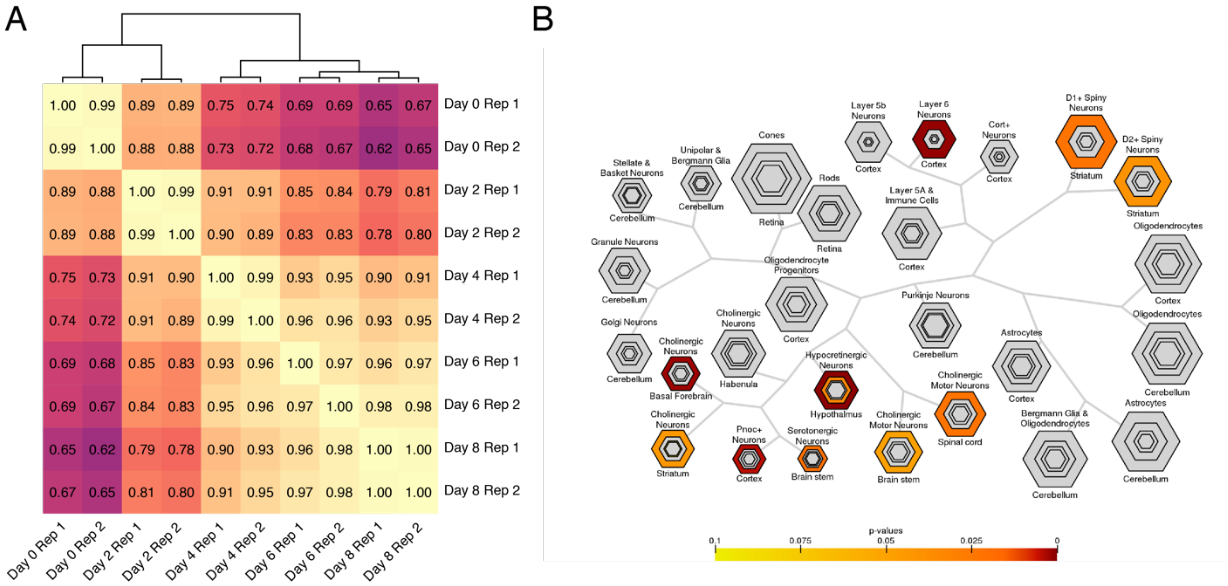


**Supplementary Figure 1: Reproducibility and cell types of LUHMES differentiation. A)** Pearson correlation of RNA-sequencing counts between all pairs of RNA-seq samples shows high reproducibility of differentiation. Samples cluster in progressive temporal order by day of differentiation. Hierarchical clustering indicates Day 4-8 samples are more related to each other than to early neuronal progenitor cells (Day 0 and Day 2 samples). n = 2 replicates for each timepoint. **B)** Cell-type specific enrichment analysis of Day 8 differentiated LUHMES shows transcriptional similarities to a variety of neurological disease relevant neuronal cell types.

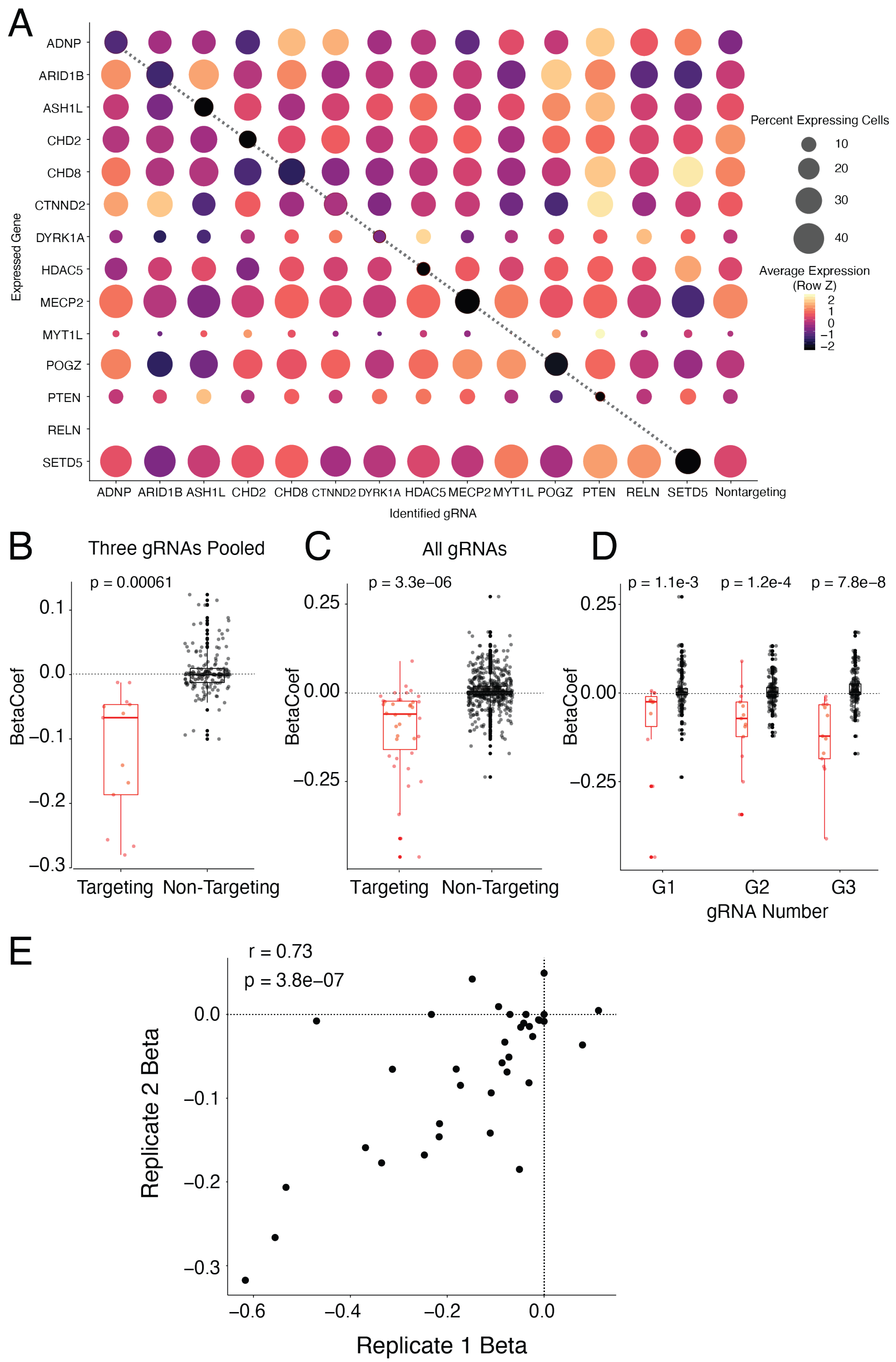


**Supplementary Figure 2: MIMOSCA analysis confirms gRNA knock-down efficiency is high and reproducible across all targeted genes. A)** Seurat DotPlot representation of the expression of ASD candidate genes (rows) across cells grouped by targeted gene (columns). Expression values are Z-normalized within a row across cells grouped by targeted gene. Cooler colors indicating lower expression. The dark diagonal reflects efficient repression in cells with a guide targeting a given gene. *RELN* was not appreciably detected in scRNA-seq data. **B)** MIMOSCA beta coefficients for targeting and non-targeting guides. Beta < 0, marked with dotted line, represents repression. Collectively, targeting gRNAs have significant, on-target activity (negative beta coefficients) when all 3 gRNAs are merged or **C)** when gRNAs are analyzed separately. **D)** All 3 designed gRNAs for each gene have significant on-target repression. Beta coefficients were not significantly different by guide number. **E**) MIMOSCA Beta estimates for individual gRNA knock-down efficiencies are highly reproducible across replicate experiments.

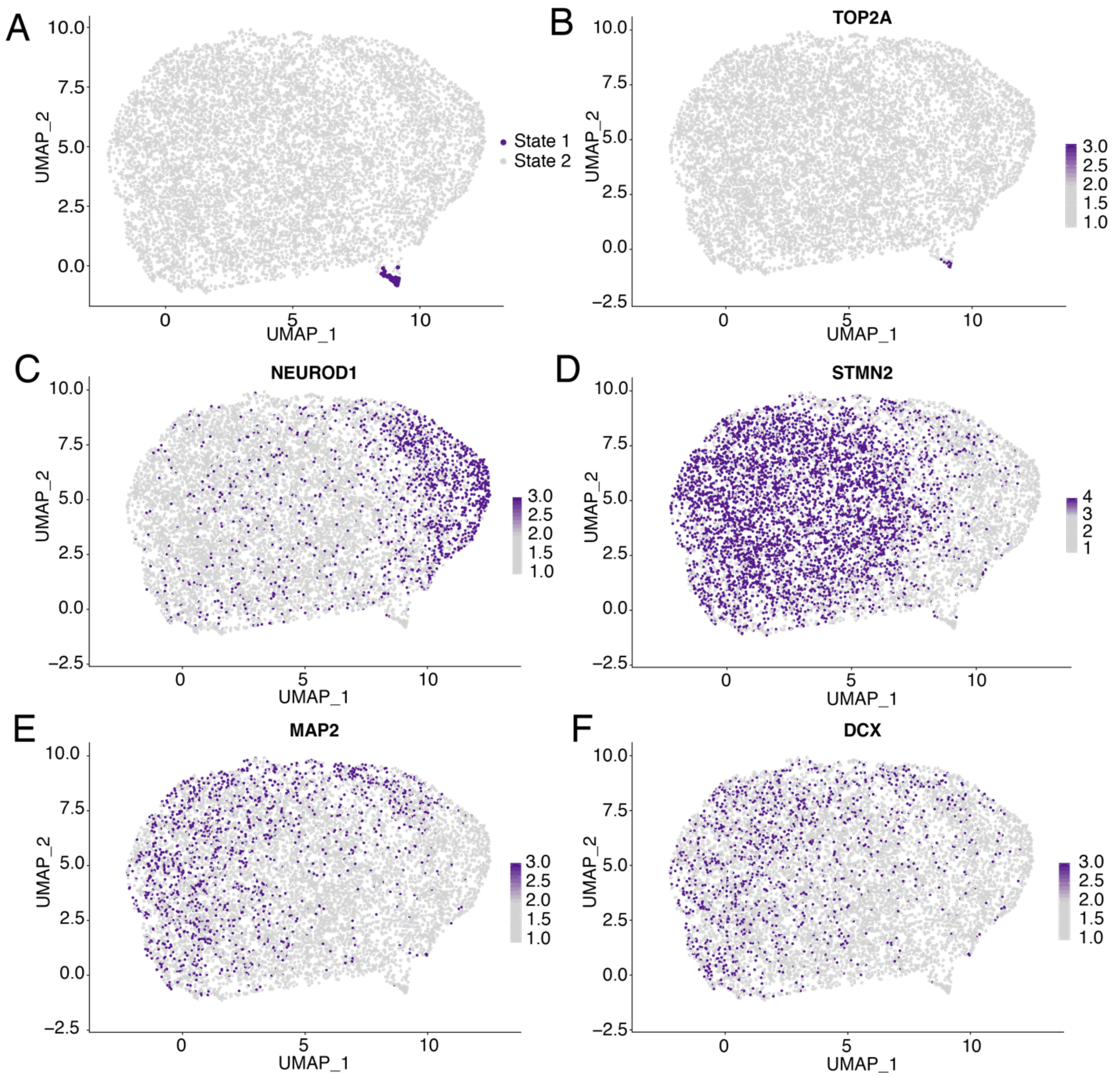


**Supplementary Figure 3: Global structure in transcriptional data.** A) UMAP clustering of transcriptomes reveals minor global variation and two cell states. B) Cell State 1 (45 cells) is enriched for expression of the DNA-replication associated gene TOP2A, a marker of proliferation; hence virtually all cells are post-mitotic. **C-F)** Neuronal marker genes show variable patterns of gene expression.

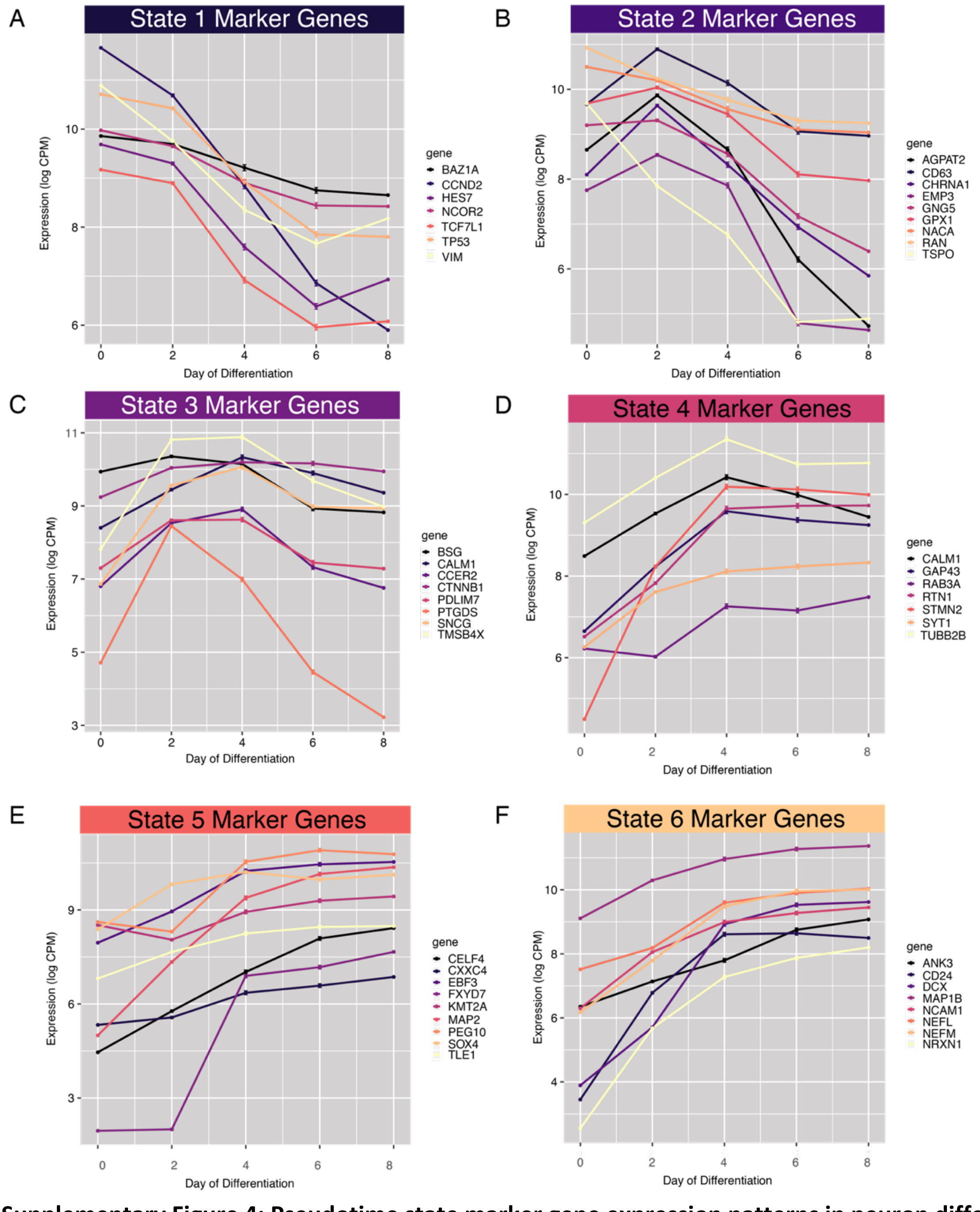


**Supplementary Figure 4: Pseudotime state marker gene expression patterns in neuron differentiation time-course dataset show pseudotime states correspond to day of differentiation.** Expression of positive marker genes from each of the six pseudo-time states in bulk RNA-seq differentiation dataset. **A)** State 1 marker genes tend to have high expression in differentiation day 0-2 cells, and include cell cycle genes (*CCND2),* proliferation genes (*TP53),* and the neural stem cell marker vimentin *(VIM)***. B)** State 2 marker genes have highest expression in day 2-4 cells**. C)** State 3 marker also peak in differentiation days 2-4. **D-F)** States4-6 express marker genes that have high expression in differentiation days 6-8. **F**) State 6 Marker genes include the mature neuronal markers *DCX, MAP1B,*

*NCAM1,* and *NEFL.* Together, these data support the interpretation of pseudo-time as an axis of progressive differentiation.

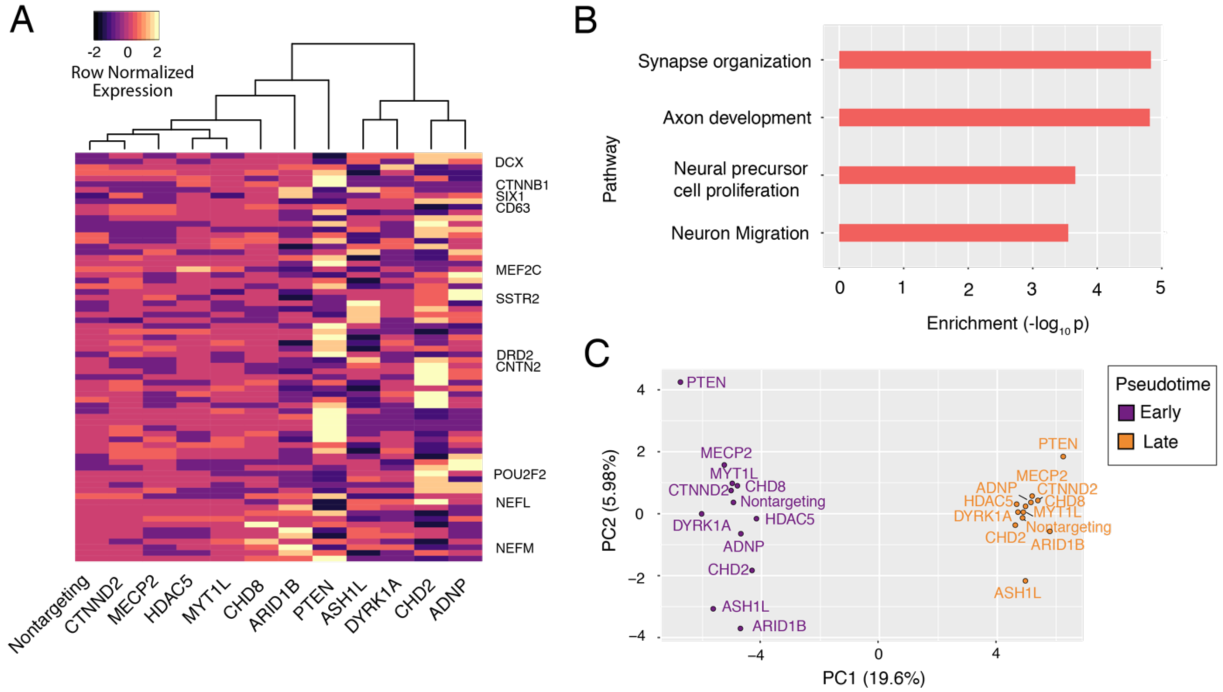


**Supplementary Figure 5: Single-cell differential expression analysis reveals variation in neuron differentiation genes. A)** Hierarchical clustering of single-cell expression of recurrently dysregulated genes. Each column represents the average expression profile of all cells having any of the 3 guides targeting the listed gene. Each row is a gene that was found to be differentially expressed in at least 3 knock-down conditions. These genes include the neuronal maturation markers *DCX, NEFL,* and *NEFM.* The 'Delayed Differentiation' module uncovered by pseudotime analysis clusters together and shows a decrease of differentiation markers. *PTEN* shows the opposite expression pattern. **B)** Biological process enrichment of recurrently dysregulated genes highlights variation in neuronal maturation processes, and neural precursor cell proliferation. **C)** Principal component analysis after stratifying expression profiles by pseudotime status shows that PC1 corresponds to pseudotime and explains 19.6% of the variance in the data. PC2 may correspond to early-stage effects of specific ASD-gene repression, and clusters the 'Delayed Differentiation' module genes together, with *PTEN* on the opposite spectrum.

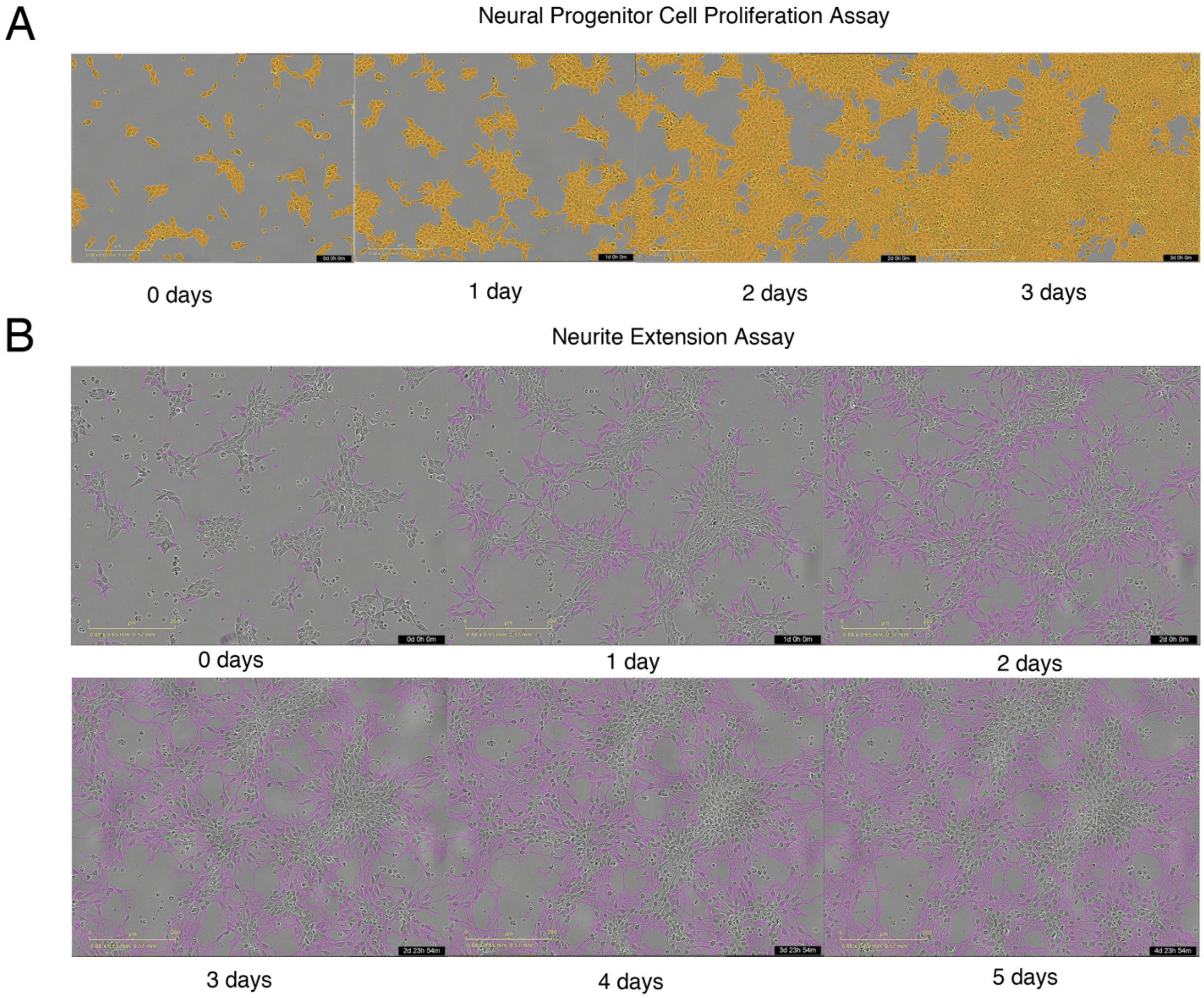


**Supplementary Figure 6: Time-lapse imaging of neural progenitor cell proliferation and neurite extension. A)** Neural progenitor cell proliferation is measured by creating a cell mask (orange) and computing the area of confluence at each time point. **B)** Neurite extension is measured with the NeuroTrack assay in the IncuCyte software. Neurite masks are shown (purple). Neurite extension lengths are normalized by cell cluster area. For both assays, images were acquired every 4 hours. Proliferation assay was performed for 3 days. Neurite extension was monitored for 5 days.

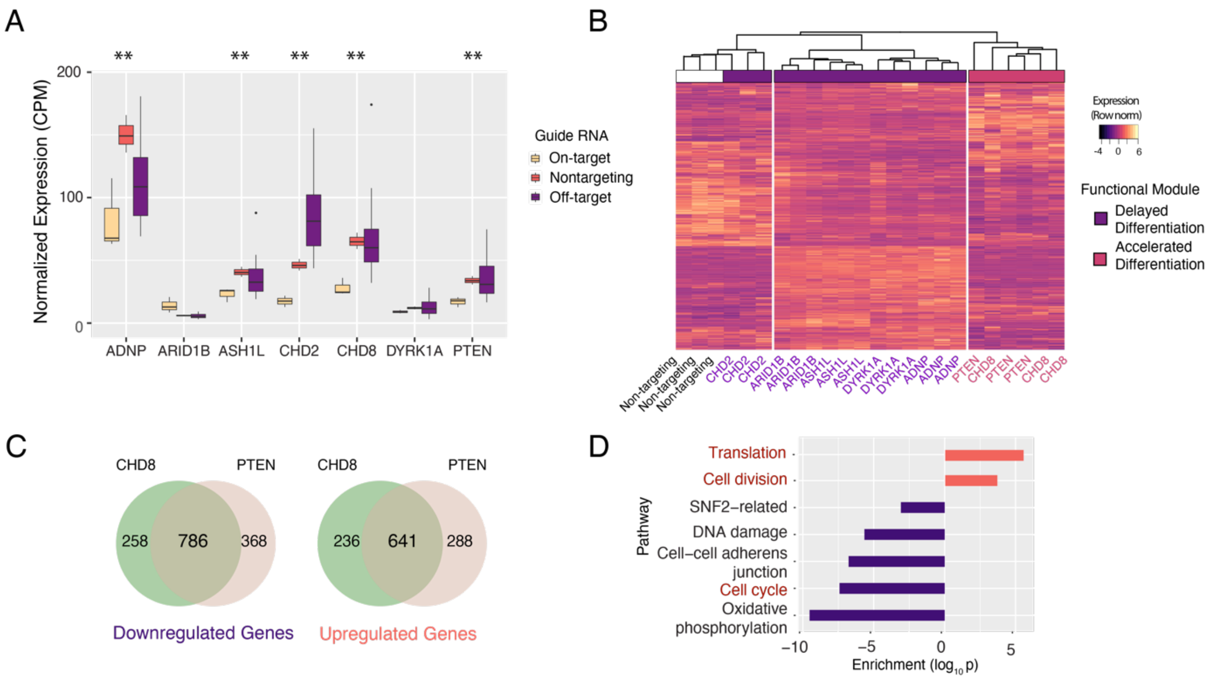


**Supplementary Figure 7: ASD-gene repression in iPSC-derived neural progenitor cells confirms module gene membership and transcriptional convergence. A)** Gene expression level in RNA-seq data confirms efficient repression for 5/7 targeted genes. Low detection of *ARID1B* and *DYRK1A* precludes interpretation. (** = $p$ < 1e-5). **B)** Unsupervised, hierarchical clustering of RNA-seq replicates by highly variable genes clusters functional module genes together. **C**) *CHD8* and *PTEN* repression elicits shared transcriptional consequences on down- and up-regulated genes. **D)** These genes are enriched for increased cell division, translation, and decreased cell-cycle genes. Downregulated genes are plotted with negative enrichment scores.

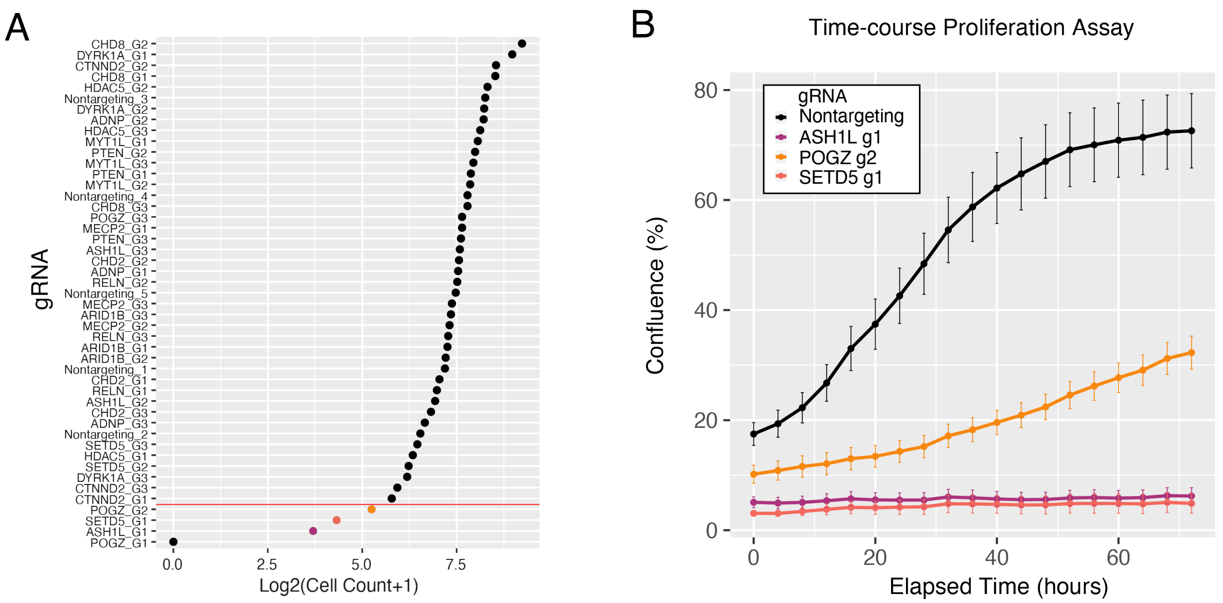


**Supplementary Figure 8: gRNA abundance in final single-cell dataset. A)** Four gRNAs were significantly depleted from the pooled single-cell experiment (chi-squared test, $p < 0.01$). **B)** Proliferation assay in LUHMES infected with individual depleted gRNAs reveals severe reductions in neural progenitor cell proliferation.

**Supplementary Figure 9: Pseudotime States Capture Global Transcriptional Differences and Reflect Distinct Cell States.**

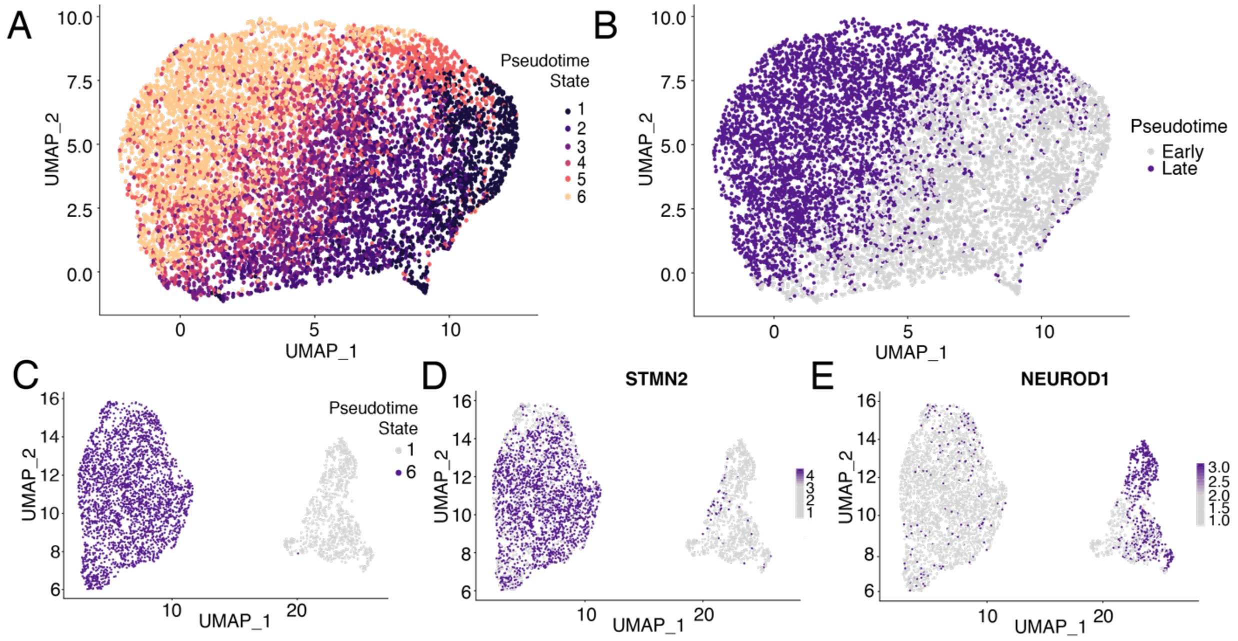


**A)** Pseudotime states labels (see Figure 3A) are transferred to UMAP. **B)** Early and Late Pseudotime State labels bisect the UMAP plot. **C)** Clustering only cells in Pseudotime states 1 and 6 reveals two discrete neuronal populations. **D)** The left cluster expresses later neuronal marker *STMN2*. **E)** The right cluster expresses early neuronal transcription factor *NEUROD1.*
